# Acute dietary methionine restriction highlights sensitivity of neocortex development to metabolic variations

**DOI:** 10.1101/2024.09.10.612174

**Authors:** Sulov Saha, Clémence Debacq, Christophe Audouard, Thomas Jungas, Pierrick Dupre, Mohamad-Ali Fawal, Clément Chapat, Henri-Alexandre Michaud, Laurent Le Cam, Matthieu Lacroix, David Ohayon, Alice Davy

## Abstract

Methionine -an essential amino acid that has to be provided by nutrition- and its metabolite S-Adenosyl methionine (SAM) are indispensable for cell proliferation, stem cell maintenance and epigenetic regulation ^1–5^, three processes that are central to embryonic development ^6^. Previous studies using chronic dietary restriction of methyl donors prior to and during gestation indicated that methionine restriction (MR) is detrimental to the development or growth of the neocortex ^7,8^, however, the consequences of acute MR have not been extensively studied. Here, we designed a dietary MR regime coinciding with the neurogenic phases of neocortex development in the mouse. Our results indicate that dietary MR for 5 days leads to a severe reduction in neocortex growth and neuronal production. In comparison, growth of the liver and heart was unaffected, highlighting an organ-specific response to MR which was also observed at the cellular and molecular levels. Progenitor cohort labeling revealed a time-dependent sensitivity to MR and cell cycle analyses indicated that after 5 days of MR, progenitors are stalled in the S/G2 phases. Unexpectedly, neocortex growth reduction induced after 5 days of MR is completely rescued at birth when switching the dam back to control diet for the remaining of gestation, uncovering a mechanism of catch-up growth. Using multiplexed imaging we probed metabolic and epigenetic markers following MR and during catch-up growth and show that pyruvate metabolism is rewired in progenitors. Altogether, our data uncover a transient state of quiescence in G2/S which is metabolically distinct from G0 quiescence and associated with efficient catch-up growth. More globally, our study highlights both the extreme sensitivity of the developing neocortex to acute dietary changes and its remarkable plasticity.

## MAIN

Methionine, an essential amino acid in mammals, is required for a number of key cellular functions, including translation, proliferation and epigenetic regulation. As a consequence, the responses to methionine restriction (MR) are pleiotropic but converge at the cellular level on an evolutionary conserved cell cycle arrest ^9^ whose underlying molecular mechanism is not completely understood. In pluripotent stem cells, MR promotes exit of the cell cycle and differentiation ^2^ and several studies have shown that methionine metabolism controls pluripotency and stem cell self-renewal by sustaining methylation reactions ^1–4^. The latter is explained by the fact that the first metabolite produced by 1C metabolic conversion of methionine is the universal methyl donor S-Adenosyl methionine (SAM) (Fig.1a), which directly links methionine levels to epigenetic regulation ^10^. Reduction in the levels of the methyl donor SAM following MR is also responsible for triggering cell cycle arrest ^11^. Methionine being provided by nutrition in mammals, we imposed MR by dietary intervention. We used modified chow that contains no methionine and normal ranges of folate and choline levels (∼2mg/kg and ∼2500mg/kg) (Extended data Table1) and first validated that intake of control and MR chow by virgin adult females and pregnant dams was equivalent, thus avoiding confounding effects of general under nutrition (Extended data Fig1a-c). To assess the consequences of acute dietary MR on neocortex growth and neuronal production, we designed dietary interventions specifically targeting the neurogenic phase while sparing earlier phases of embryonic development. Pregnant females were subjected to MR starting at E9.5 for 5 days until E14.5 or for 10 days until E19.5/P0 (Fig1b). Dietary MR for 10 days led to intrauterine growth restriction (IUGR) shown by decreased body weight (Fig1c) and a survey of different organs indicated that brain weight was decreased but not liver and heart weights (Fig1c). Next, we analyzed the consequences of dietary MR for 5 days. At E14.5, no IUGR could be detected, yet a decrease in brain weight was already present (Fig1d), while no difference in liver or heart weight could be detected (Fig1d). To assess whether dietary MR led to organ-specific changes in methionine levels, we measured steady-state methionine levels and observed that 5 days MR led to a decrease in methionine levels in the neocortex and in the liver but not the heart of E14.5 embryos (Fig1e). Altogether, these results reveal an organ-specific response to dietary MR and indicates that the brain is more sensitive to MR than other organs.

To explore the molecular and cellular responses to decreased methionine levels, we quantified global DNA methylation by ELISA (enzyme-linked immunosorbent assay) and assessed cell cycle parameters by cytometry. We observed a significant decrease in global DNA methylation in the neocortex but not in the liver or heart of E14.5 embryos subjected to dietary MR for 5 days (Fig1f). Cytometry analyses indicated that distribution in the cell cycle was modified in the neocortex in E14.5 embryos subjected to dietary MR for 5 days but not in the liver or heart (Fig1g). Specifically, in the neocortex, the fraction of cells in S and G2/M phases of the cell cycle was increased at the expense of cells in G0/G1, yet there was no increase in the proportion of cells positive for the mitotic marker phospho-Histone H3 (pH3) (Fig1g). These data indicate that neocortical progenitors are arrested or delayed in the S and G2 phases of the cell cycle in response to dietary MR for 5 days.

Development of the neocortex is a stepwise process that involves two broad types of progenitors in the developing cortex, apical progenitors that express the transcription factor PAX6 and intermediate basal progenitors that express the transcription factor TBR2 ^12–14^. Collectively, these progenitors give rise to two broad categories of neurons, early born neurons that express the transcription factor CTIP2 and late born neurons (which start to be produced at E14.5) expressing the DNA binding protein SATB2. These different cell types are organized in different layers in the developing tissue: apical progenitors are located in the ventricular zone (VZ), newborn neurons and intermediate progenitors form the intermediary zone (IZ), while differentiated neurons are grouped in the cortical plate (CP) (Fig2e, i). At later stages, a fraction of progenitors give rise to glial cells (astrocytes and oligodendrocytes) ^15^ (Fig3g). Proliferation, self-renewal/differentiation and epigenetic remodeling must be coordinated throughout this developmental process to ensure proper neuronal and glial production both in term of number and subtypes ^16^.

We characterized the consequences of the dietary interventions described above (Fig2a) both on neocortex growth and on neuronal production. As shown above, MR for 10 days led to a reduction in brain weight observable macroscopically (Fig2b). We quantified this reduction in neocortex growth by measuring radial thickness and tangential extension on coronal sections (Fig2c). After 5 days of MR, radial thickness was already reduced (Fig2d), however, decreased radial thickness was not observed after 3 days of MR (Extended data FigS2a-c). To assess whether the reduction in growth following 10 days of MR correlates with the loss of specific neuronal populations, we performed immunostaining with markers of newborn neurons (TBR1), deep layer neurons (CTIP2) or late born neurons (SATB2) (Fig2e-h). Quantification of the different markers revealed that MR for 10 days leads to a global reduction of all neuronal subtypes in the neocortex (Fig2h). After 5 days of MR, the number of all neuronal subtypes was already strongly reduced (Fig2i-l) and this was not due to increased cell death (Extended data FigS2d, e). We then quantified progenitor numbers using markers of apical progenitors (PAX6) and intermediate basal progenitors (TBR2) (Fig2m). Quantification of the two markers indicated that both populations of progenitors are decreased after 5 days of MR (Fig2n). The cycling properties of these progenitors were analyzed by EdU incorporation which revealed an increased proportion of EdU+ cells following 5 days of MR, as well as an increased proportion of TBR2+/EdU+ intermediate progenitors (Fig2o-p). Conversely, the proportion of pH3+/EdU+ mitotic cells was decreased in MR conditions indicating that EdU+ progenitors do not proceed to mitosis, thus supporting the interpretation that they are stalled in the S or G2 phases of the cell cycle. Altogether these results indicate that dietary MR leads to severe neocortex growth defects and microcephaly within 5 days, by altering the cycling properties of neural progenitors.

It has been shown previously that neuronal production in the developing neocortex is a plastic process that can adapt to a transient loss of cells or to a lower rate of differentiation ^17–19^. To assess whether restricted growth of the neocortex induced by 5 days of MR is reversible, we designed a regimen in which after 5 days of MR, the pregnant dams were switched back to control diet (Fig3a) and brain and neocortex growth were assessed at E19.5/P0. With this regimen, no gross difference in brain size was detected at E19.5/P0 (Fig3b), tangential growth of the neocortex was rescued (Fig3c) and radial growth of the tissue was less severely decreased compared to 10 days of MR. No difference in neuron numbers was observed between control regimen and the switched MR regimen (Fig3d-f), indicating that all three neuronal subtypes were completely rescued despite the severe microcephaly that was present at E14.5. Next, we assessed whether this rescued neuronal production was done at the expense of glial production which in normal conditions starts around E16.5. To do this, we exposed Aldh1L1-EGFP reporter mice to the three diet regimen (Fig3g) and assessed TBR2 as a marker of the neuronal lineage and used ALDH1-L1, OLIG2 and SOX9 as markers of glial cells at E19.5. As expected, following 10 days of MR we observed a decrease in the total number of cells (DAPI+) which was compensated in the switched condition (Fig3h-i). No statistically significant difference in the number of TBR2+ cells was observed following 10 days of MR while the number of glial cells (ALDH1-L1+, OLIG2+, SOX9+) was decreased (Fig3j-k). Interestingly, in the switched diet condition, the number of SOX9+ glial cells was not completely compensated (Fig3j-k) indicating that glial cell production was hampered or delayed during catch-up growth. These results indicate that MR-induced cell cycle stalling is reversible and they reveal a mechanism of catch-up growth in the developing neocortex which compensates neuronal numbers at the expense of glial cell production.

In order to better understand progenitor behavior following MR and during catch-up growth, we used cell cohort tracing approaches in combination with diet regimen (Fig4a). We labeled a cohort of neural progenitors at E12.5 with EdU (labeling neural progenitors in S-phase) and in the same pregnant dam, we labeled a cohort of neural progenitors at E14.5 using in utero injections of FlashTag/CFSE (labeling mainly mitotic neural progenitors abutting the ventricle) ^20,21^. In this set of experiments, the switch between MR and control diet was done at E14, 15 hours before FlashTag injection, in order to label progenitors in the catch-up growth phase (Fig4a). Injected embryos were then analyzed by immunofluorescence 2 days later, at E16.5 (Fig4b). The proportion of EdU+ and FlashTag (FT)+ cells was quantified in the different cell layers of the neocortex (ventricular zone (VZ), intermediary zone (IZ), cortical plate (CP)). Focusing on EdU+ cells, the quantification at E16.5 shows that in all dietary conditions, the majority of neural progenitors that were cycling at E12.5 are located in the deep layer of the CP, suggesting that they differentiated into deep layer neurons. No significant difference was observed between the three diet conditions (Fig4c). Focusing on FT+ cells, the quantification at E16.5 shows that in the control situation, a large fraction of neural progenitors that were undergoing mitosis at E14.5 have differentiated, and are already positioned in the upper layers of the CP while some FT+ cells are located in the IZ (Fig4d). Very few FT+ cells have remained in the VZ. Following 7 days of MR, a large proportion of FT+ cells are still located in the VZ and IZ with few having reached the CP (Fig4d) indicating defects and/or delay in differentiation or migration. In the switched regimen, while many FT+ cells are present in the VZ and IZ, a large proportion of them has reached the CP (Fig4d), indicating that switching back to control diet has allowed FT+ cells to quickly resume differentiation and migration.

Altogether these results suggest that MR induces a reversible quiescent-like state in mid-to-late progenitors which would be quickly exited upon favorable dietary conditions. Some of the hallmarks of quiescence are downregulation of Ki67, modifications of epigenetic landscape and energy metabolism as well as a shutdown of translation ^22–24^. To further characterize the quiescent state of progenitors following MR and during catch-up growth we performed imaging mass cytometry that allows to interrogate multiple proteins and protein posttranslational modifications simultaneously ^25,26^. We used a panel of antibodies (Extended data Table 2) including enzymes involved in glycolysis and in the TCA cycle, a marker of translation (4E-BP1), Ki67 as a marker of proliferating cells and the progenitor marker Vimentin. Since it has recently been shown that proliferating progenitors produce high levels of lactate that promote blood vessel formation in the developing neocortex ^27^, we also tested markers of blood vessels and oxygenation (CD31, CA9, GLUT1). Lastly, we included antibodies specific for histone modifications (methylation and acetylation) to assess epigenetic marks. Of note, IMC and image analyses were performed on two sets of samples. For clarity, data from only one set are presented in the main figure while selected data from the second set are presented in Extended data FigS3. Sections of the neocortex of E16.5 embryos subjected to the 3 dietary regimens described above were processed for IMC and multiplexed pseudo-colored images were reconstituted to visualize the expression of different markers focusing on the ventricular zone (VZ) (Fig4e). Images were segmented (Fig4f) and bioinformatic analysis of the IMC data was done using HistoCAT ^28^. Unsuperviszed non-linear dimentional reduction using t-distributed stochastic neighbor embedding (t-SNE) revealed three clusters corresponding to the three different diet conditions (Fig4g and Extended data FigS3), indicating that the three diet conditions exhibit distinct profiles of marker expression. To further analyze the IMC data we used QuPath ^29^ to quantify the fraction of nuclei positive for a given marker as described by others ^30^. This analysis indicates that the fraction of Ki67+ cells in the VZ was increased in 7 days MR condition compared to control or switched diet conditions (Fig4h and Extended data FigS3). This is consistent with the fact that progenitors are stalled in S or G2 phases of the cell cycle and not in G0. Further, an increase in 4E-BP1 levels was observed following 7 days MR compared to control or switched diet conditions (Fig4h and Extended data FigS3), indicating that translation is inhibited following MR and restored in the switched diet condition. No difference in blood vessels or oxygenation markers was observed. Concerning histone methylation and acetylation marks, the main difference observed was a reduction in the fraction of cells positive for H3K27me3 following MR and in the switched diet condition (Fig4h). Metabolic markers that were tested by IMC participate in pyruvate metabolism or the TCA cycle and OXPHOS (Fig4i). Since these metabolic markers are not enriched in the nucleus, we used HistoCAT to quantify mean signal intensity in the three diet conditions. Interestingly, for the majority of metabolic markers, we observed an increase in the mean intensity signal in 7 days MR compared to control or switched diet conditions (Fig4j and Extended data FigS3). Two notable exceptions are p-PDHA1 and PC which were decreased in 7 days MR. Collectively, these changes suggest that progenitors subjected to 7 days MR exhibit modified pyruvate metabolism compared to control diet or to catch-up growth conditions. Altogether, these data reveal that following MR, progenitors exhibit hallmarks of quiescence including a shift in energy metabolism, changes in epigenetic marks and inhibition of translation.

Our study reports that the response to dietary MR is fast, with a severe, yet reversible, deficit in neuronal and glial production that correlates with rewiring of energy metabolism in progenitors. Following MR, neural progenitors are stalled in the S or G2 phases of the cell cycle leading to a general decrease in the number of cells produced (progenitors, neurons and glial cells). While an MR-induced cell cycle block in S or G2 has been reported in prostate cancer cells ^31^, in many other cancer cells that are highly dependent on external methionine, an arrest in G0/G1 is observed following MR ^32^. Interestingly, a G2-quiescence state was identified in Drosophila neural progenitors and it was shown that progenitors in G2-quiescence reactivate faster than progenitors in G0-quiescence in response to nutrients ^33^. In other non-mammalian contexts, G2 quiescence has been described as a shallower quiescent state allowing faster reactivation, for instance in response to injury ^34,35^. While quiescence in G0/G1 is characterized by dampening of energy metabolism, how energy metabolism is rewired in the G2-quiescent state is less understood. Our data suggest that pyruvate metabolism is modified in progenitors that are stalled in S/G2, possibly leading to increased OXPHOS. This is coherent with recent studies reporting that mitochondrial pyruvate metabolism is active in quiescent cells and that the pyruvate transporter MPC1 is needed to maintain quiescence ^36,37^.

Our study uncovers a mechanism of catch-up growth. Within 2 days of switching back to normal diet, the system has resumed and even accelerated its neuronal production at the expense of glial production. Catch-up growth is defined by the accelerated growth of a tissue to reach a certain target size appropriate for age. This concept has been largely explored with respect to bone growth ^38^. One hypothesis to explain catch-up growth is the « sizostat » mechanism which was originally formulated for whole body growth and implied a neuroendocrine loop ^39^. An alternative hypothesis proposed that the mechanism for catch-up growth was intrinsic to each tissue, involving accelerated rates of cell proliferation ^40^. Our data suggest that in the neocortex, catch-up growth may not be driven by accelerated proliferation but by a delay in the production of glial cells. Altogether, our study emphasizes the metabolic vulnerability and plasticity of the developing neocortex.

## METHODS

### Animals

Wild type mice were kept in a 129S4/C57Bl6J mixed background. Aldh1L1-EGFP ^41^ mice were from the GENSAT project and genotyped as shown previously ^42^. The sex of embryos was not tested and littermates were randomly assigned to replicate experiments. Mice were maintained on a standard 12 hours day:night cycle and given ad libitum access to food and water. Mice were either fed a methionine-restricted diet (SAFE AIN 2g Choline #U8958 Version 330) or a control diet (SAFE #AO4) during pregnancy. Timed pregnancies were obtained by detecting vaginal plugs in the morning and samples were collected in the afternoon. All experimental procedures were pre-approved and carried out in compliance with the guidelines provided by the national Animal Care and Ethics Committee (APAFIS #35692-2022082409293050 v6) following Directive 2010/63/EU.

### Evaluation of food intake

To measure food intake, a group of 2-3 non-pregnant or pregnant females (3-5 months old) were kept in a cage. A defined amount of food was provided to the mice, the remaining food was weighed at 2 days interval for the calculation of average food intake.

### Weight measurement

Body weight was measured after the removal of placenta. The wet weight of fresh organ samples was recorded as soon as it is removed from the body cavity.

### Immunohistochemistry

Mouse brains at stages E14.5 and older were collected and fixed with 4% Paraformaldehyde overnight at 4°C followed by paraffin or OCT embedding. The paraffin embedding was obtained with Spin tissue processor (Myr) for 5 µm coronal sections on Microtome (Microm). For OCT embedding, the samples were transferred to 30% Sucrose solution overnight at 4°C before OCT (Sakura) inclusion for performing 20 µm coronal sections using Cryostat (Leica Biosystems). Cryostat sections were washed three times with PBST (1x PBS with Tween 20) and then incubated for 1 hour at room temperature with blocking buffer (4% goat serum in PBST). Then, sections were incubated with antibodies diluted in blocking buffer overnight at 4°C. Paraffin-embedded sections were exclusively deparaffined, rehydrated, followed by antigen retrieval involving steam heating for 45 minutes. Coronal sections were permeabilized and incubated in blocking solution (1-3% BSA, 1-3% FBS, 0.5-1% Tween 20 and PBS) for 2 hours at room temperature. Sections were permeabilized and incubated with antibodies diluted in blocking solution (1-3% BSA, 1-3% FBS, 0.5-1% Tween 20 and PBS) overnight at 4°C.

For birthdating experiments, EdU (dissolved in PBS at 5 mg/ml, 25 μg/g body weight) was injected intraperitoneally at E12.5. Histological detection of EdU was performed using the Click-iT EdU cell proliferation kit (ThermoFisher) according to the manufacturer’s instructions. A stock of 10mM FlashTag (CFDA-SE, STEMCELL Technologies) was prepared by dissolving the CFDA-SE in dimethyl sulfoxide (DMSO) (Euromedex) for an in utero injection. In utero injections were performed as described previously ^43^. The fresh working solution of 1mM was diluted in HEPES-buffered saline solution (HBSS, ThermoFisher), in which Fast Green dye was added for monitoring successful injection.

### Imaging mass cytometry

Mouse brains at E16.5 were collected and fixed with 4% Paraformaldehyde overnight at 4°C followed by paraffin embedding. A 3-μm-thick coronal section of the brains was processed in a workflow analogous to immunohistochemistry slide preparation. A region of interest, ROI (dorsolateral neocortex) in different conditions was identified in hematoxylin and eosin-stained sections. Tissue sections were prepared for the deparaffination and antigen retrieval steps using Dako target retrieval solution pH9 (Agilent) at 96°C during 30 minutes. After that, nonspecific binding sites were blocked with blocking buffer (Mouse FcR blocking reagent, Miltenyi) diluted at 1:100 within SuperBlock blocking buffer (ThermoFisher) and then stained overnight at 4°C with antibody mix diluted in PBS-1%BSA (BSA Cohn fraction 30%, Sigma-Aldrich). After three 8 minutes washing steps (Triton X-100 diluted at 0.2% within dPBS), the slide was stained with a nuclear intercalator solution containing metal isotopes (Intercalator-Iridium diluted at 1:500 within dPBS). The slide was then washed 5 minutes in ultrapure water and left to dry before IMC acquisition. All antibodies (see Extended data Table 2) were conjugated to metals using the MaxPar X8 multimetal labeling kit (Standard BioTools) according to the manufacturer’s instructions. Previously tested antibodies were chosen for metal conjugation and targets of interest. Data acquisition was performed on a Helios time-of-flight mass cytometer (CyTOF) coupled to a Hyperion imaging system (Standard BioTools). Selected areas for ablation were larger than the actual area of interest to account for loss of overlapping areas among sections due to cumulative rotation. The selected area ablated per section was around 1 mm^2^. Laser ablation was performed at a resolution of approximately 1 µm with a frequency of 200 Hz. All IMC data from two batches were stored as MCD files.

### Imaging and quantification

The macroscopic images were acquired with SMZ18 stereomicroscope (Nikon) and fluorescent images were captured with SP8 inverted or upright scanning confocal microscope (Leica Biosystems). Images were exported as TIFF files and manual and semi-automated quantification was performed with ImageJ (NIH) and QuPath scripts on defined regions of interest (ROIs)/hemisphere. Embryos were collected from different litters and cells were counted per ROI/hemisphere using multiple sections (see Extended data Table 3 for raw counts). For E12.5 mouse embryos, one ROI in the dorsolateral region was chosen for manual counting. 100-200 cells were counted per ROI/hemisphere and counts were normalized to an area of 100 μm of apical surface×100 μm of radial length. For E14.5 and E16.5 mouse embryos, one ROI in the dorsolateral region/whole hemisphere was chosen for semi-automated segmentation and quantification using “NucleusDetection” script in the QuPath software. We manually defined a threshold to identify DAPI-stained nuclei and create a mask of all nuclei. Positive nuclei for different markers were automatically quantified using pixel intensity of fluorescent channels. For E19.5/P0 mouse embryos, a custom script was developed for semi-automated cell segmentation and quantification in ImageJ. We defined a threshold to identify nuclei in the DAPI channel and create a mask of all nuclei. Using image filtering on fluorescent channels, we further considered two parameters (>50 μm2 area of the fluorescence signal and >0.6 circularity index) to identify the positive nuclei across the entire neocortical hemisphere. All semi-automated quantification methods were validated by carrying out double-blind comparisons of manual and semi-automated quantification on the same samples. For Aldh1L1-EGFP transgenic mouse embryos at E19.5/P0, one ROI (dorsolateral neocortex) in the hemisphere was chosen for semi-automated segmentation and DAPI quantification using “NucleusDetection” script on the QuPath software. SOX9, OLIG2 and EGFP-positive cells were manually counted per 500 μm of apical surface×respective radial length.

For imaging mass cytometry, some markers were eliminated from the analysis based on these reconstituted images (unexpected pattern or absence of signal) (Extended Table 2). We used a similar pipeline to segment the images with QuPath. Segmentation was done on nuclei and the mask were expanded to include a cytoplasmic ring. Cells were classified by quantifying gray values (ranging from 0 to 255) and setting minimum thresholds based on the detection of Histone H3 and H4 signals. For each marker, cells with a signal above the threshold were classified as positive. For low-abundant proteins, the threshold was further refined by incorporating puncta identification with minimum size criteria. The data is presented as the percentage of positive cells for each marker. For bioinformatic analyses, images were extracted with Steinbock ^44^, segmentation was done with Mesmer and data were exported to HistoCAT ^28^ for further analyses.

### Methionine assay

Methionine levels were determined by a fluorometric assay kit (Abcam) based on an enzymatic mechanism coupled with the generation of hydrogen peroxide. Dissected E14.5 tissue samples were mechanically dissociated with a pipette tip and suspended in 200 µl of supplied assay buffer. Liver samples were specifically pre-treated by adding 4 μl supplied Sample Clean-Up Mix in the assay buffer for 30 minutes incubation at 37°C. For neocortex and heart, cell debris were pelleted by centrifugation at 13,000 g for 15 minutes at 4°C and supernatant was collected. All samples were then filtered through a 10kD spin column (Abcam) by centrifugation at 10,000 g for 10 minutes at 4°C. In a black 96-well microplate (ThermoFisher), 20 μl of the ultra-filtrate were mixed with the assay buffer for a total volume of 50 μl sample. 10 µl of standard stock (10 mM methionine) was serially diluted with the assay buffer. 50 µl of standard mix and samples were then incubated with supplied Reaction Mix at a total volume of 100 µl at 37°C for 30 minutes in dark. The signal intensity of fluorescence excited at 535 nm for 1 second was measured at 587 nm using Varioskan flash (ThermoFisher). The readouts from standard were plotted to construct a standard curve for estimating methionine concentration.

### DNA methylation assay

DNA was extracted from 10 mg of dissected E14.5 tissue samples using the DNeasy blood & tissue kit (Qiagen) according to the manufacturer’s instructions. The amount of isolated DNA was quantified using Nanodrop spectrophotometer (ThermoFisher). 100 ng was used as input DNA for detecting global DNA methylation status by MethylFlash methylated 5mC DNA quantification kit (EpigenTek) based on an ELISA-like colorimetric reaction. The input DNA was added to supplied strip wells with high DNA affinity. The methylated fraction of DNA was detected using supplied capture and detection antibodies and then quantified from the absorbance measurement at 450 nm in Multiskan GO microplate reader (ThermoFisher). The amount of methylated DNA is proportional to the OD intensity readouts, which were used for estimating the relative methylation status by the calculation of percentage of 5-methylcytosine (5mC) in total DNA.

### Cell cycle analysis by flow cytometry

E14.5 fresh tissue samples were collected in ice-cold PBS. Heart samples were pooled from 4 embryos, all samples were transferred into individual tubes in ice-cold FCS-containing PBS (1%). The samples were further processed for cell dissociation and staining for DAPI and phospho-Histone H3 (pH3) according to the methodology described in a previous study ^45^. The tissue lysates were loaded in Cytoflex-S flow cytometer (Beckman Coulter) and data were collected and analyzed using CytExpert software (Beckman Coulter). A range of 30,000–50,000 events was acquired with a minimal flow rate (<1000 events/s) from each sample. For analysis, clumps, debris, and doublets were eliminated and positive fluorescent signal was defined with staining controls for the primary and secondary antibodies. For cell cycle analysis, DNA content histograms from PI incorporation were used for a manual determination of G1/G0, S, and G2/M phases ^46^.

### Statistics and reproducibility

For each assay/experiment, tissue samples were obtained from at least four to eight mouse embryos per condition, collected from two different litters. Imaging mass cytometry was performed on 2 embryos per conditions. For experiments involving a pair of conditions, statistical significance between the two sets of data were analyzed with unpaired t test or Mann–Whitney test with Holm-Šidák adjusted P-values using Prism10 (GraphPad). For experiments involving multiple conditions, statistical significance was analyzed using Two-way ANOVA with Tukey’s multiple comparison test using Prism10 (GraphPad). Statistically significant differences are reported on the Figures and statistical tests are indicated in the Figure legends.

## DATA AVAILABILITY

Source data for Figures 1-4 and Extended Data figures are provided with this paper in the <u>Source data</u> files. All reagents used are available from commercial sources as presented in the Extended data-Materials.

**Figure 1.**
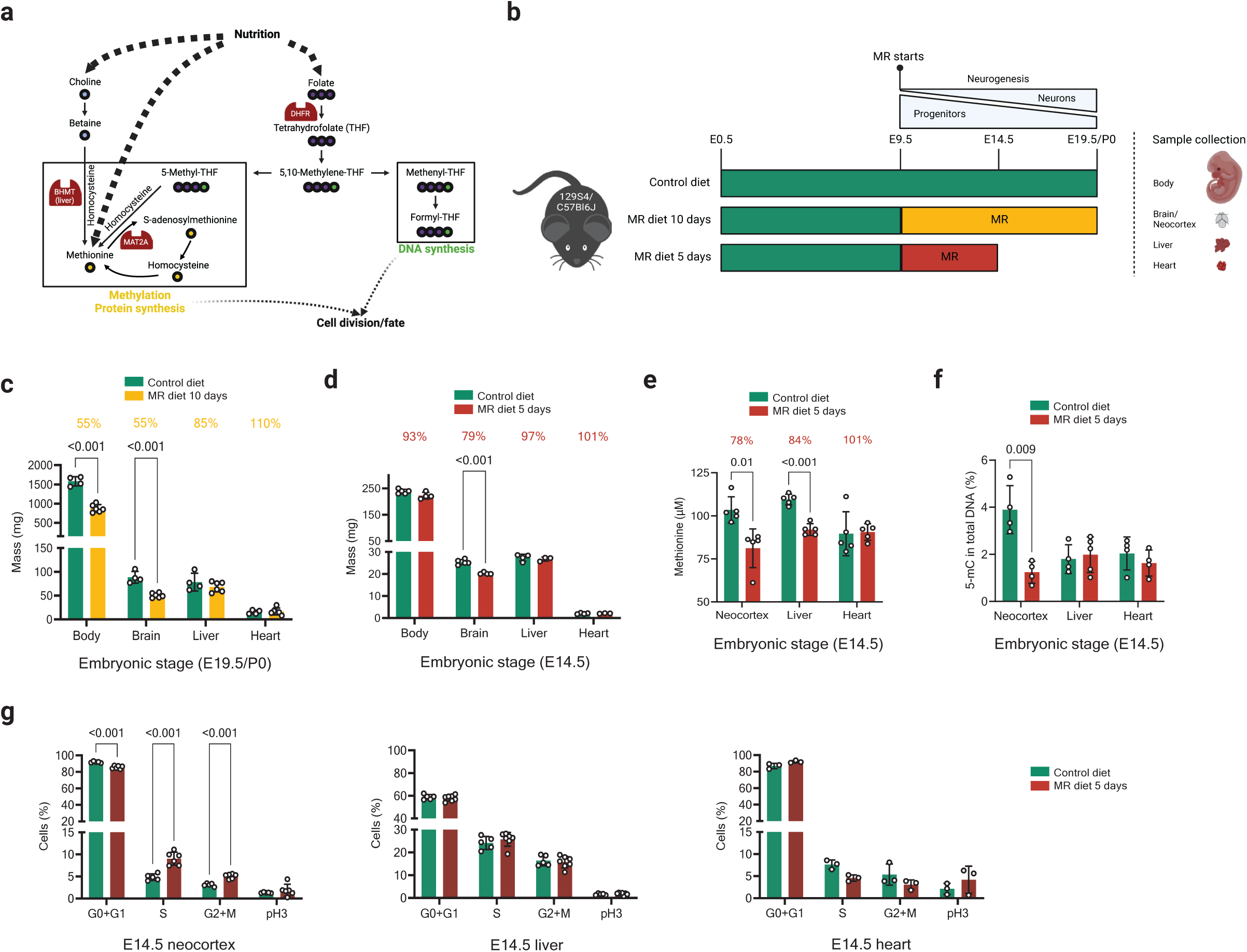
Organ-specific response to methionine restriction. **a**, Schematic representation of the 1C metabolism and its outputs. Created with BioRender.com. **b,** Schematic of study design. Pregnant dams were fed methionine restricted diet either for 5 days or for 10 days. MR was started at E9.5. **c,** Fetal body and organ weights of E19.5/P0 pups exposed to control or MR diet for 10 days (n= 4-5 embryos). **d,** Fetal body and organ weights of E14.5 embryos exposed to control or MR diet for 5 days (n= 4-5 embryos). **e,** Levels of methionine in fetal tissues extracted from E14.5 embryos exposed to control diet or 5-days MR (n= 5 embryos). **f,** Level of 5mC DNA methylation quantified in tissue extracts of E14.5 embryos exposed to control diet or 5-days MR (n= 4-5 embryos). **g,** Flow cytometry of cells dissociated from organs of E14.5 embryos exposed to control diet or 5-days MR. Cell cycle phases were identified based on DNA content (PI incorporation) and phospho-Histone H3 (pH3) staining (n= 5-7 embryos). Data are mean±s.d. Statistical analysis was carried out using multiple unpaired t test with Holm-Šídák correction. p-values are indicated when differences are statistically significant. The percentage values at the top of the graph indicate the magnitude of the mean difference between conditions.

**Figure 2.**
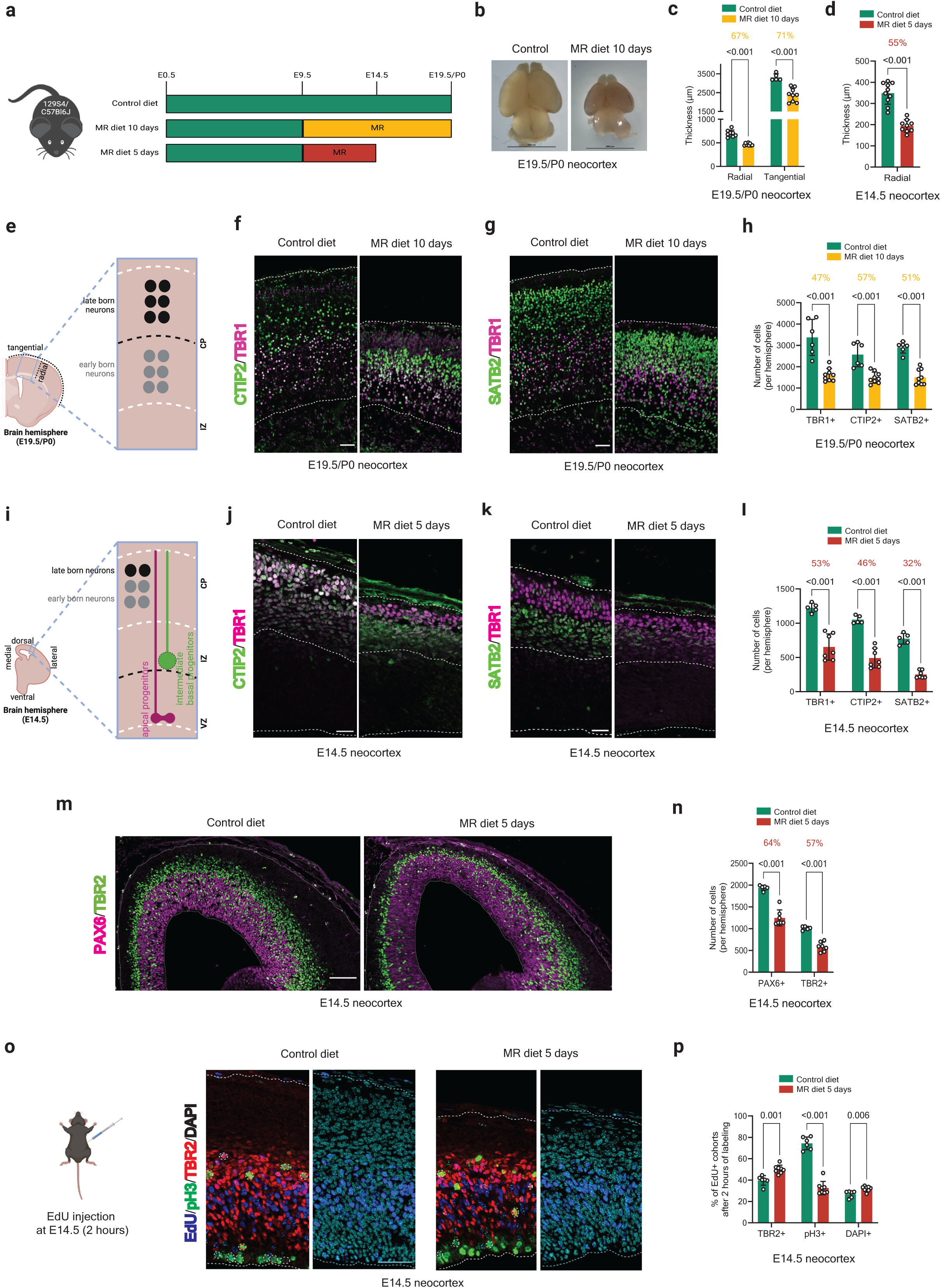
Methionine restriction leads to neocortex growth defect and microcephaly. **a**, Schematic of study design illustrating dietary regimens. **b,** Dorsal view of fetal brain at E19.5/P0 from pups exposed to control diet or 10-days MR. Scale bars: 5000 μm. **c,** Quantification of radial length and tangential perimeter of neocortex of E19.5/P0 pups exposed to control diet or 10-days MR (n= 6-8 embryos). **d,** Quantification of radial length of neocortex of embryos exposed to control diet or 5-days MR (n= 6-7 embryos). **e,** Illustration of brain hemisphere and neocortex layering at E19.5. Created with BioRender.com. **f-h,** Representative images and quantification of E19.5/P0 neocortex coronal sections immunostained for TBR1, CTIP2, and SATB2, in condition of control diet or 10-days MR. Scale bars: 50 μm. **i,** Illustration of brain hemisphere and neocortex layering at E14.5. Created with BioRender. **j-l,** Representative images and quantification of E14.5 neocortex coronal sections immunostained for TBR1, CTIP2, and SATB2, in condition of control diet or 5-days MR (n= 6-7 embryos). Scale bars: 25 μm. **m-n,** Representative images and quantification of E14.5 neocortex coronal sections immunostained for PAX6 and TBR2, in condition of control diet or 5-days MR (n= 6-7 embryos). **o-p,** Representative images and quantification of E14.5 neocortex coronal sections immunostained for EdU, phospho-Histone H3 (pH3) and TBR2, in condition of control diet or 5-days MR, at 2-hours post-EdU injection (n= 6-7 embryos). Quantification shows the percentage of cells that are EdU+ (DAPI) as well as the percentage of intermediate progenitors (TBR2+) and mitotic cells (pH3+, marked with dashed line) that are EdU+. Scale bars: 50 μm. Data are mean±s.d. Statistical analysis was carried out using multiple unpaired t test with Holm-Šídák correction. p-values are indicated when differences are statistically significant. The percentage values at the top of the graph indicate the magnitude of the mean difference between conditions.

**Figure 3.**
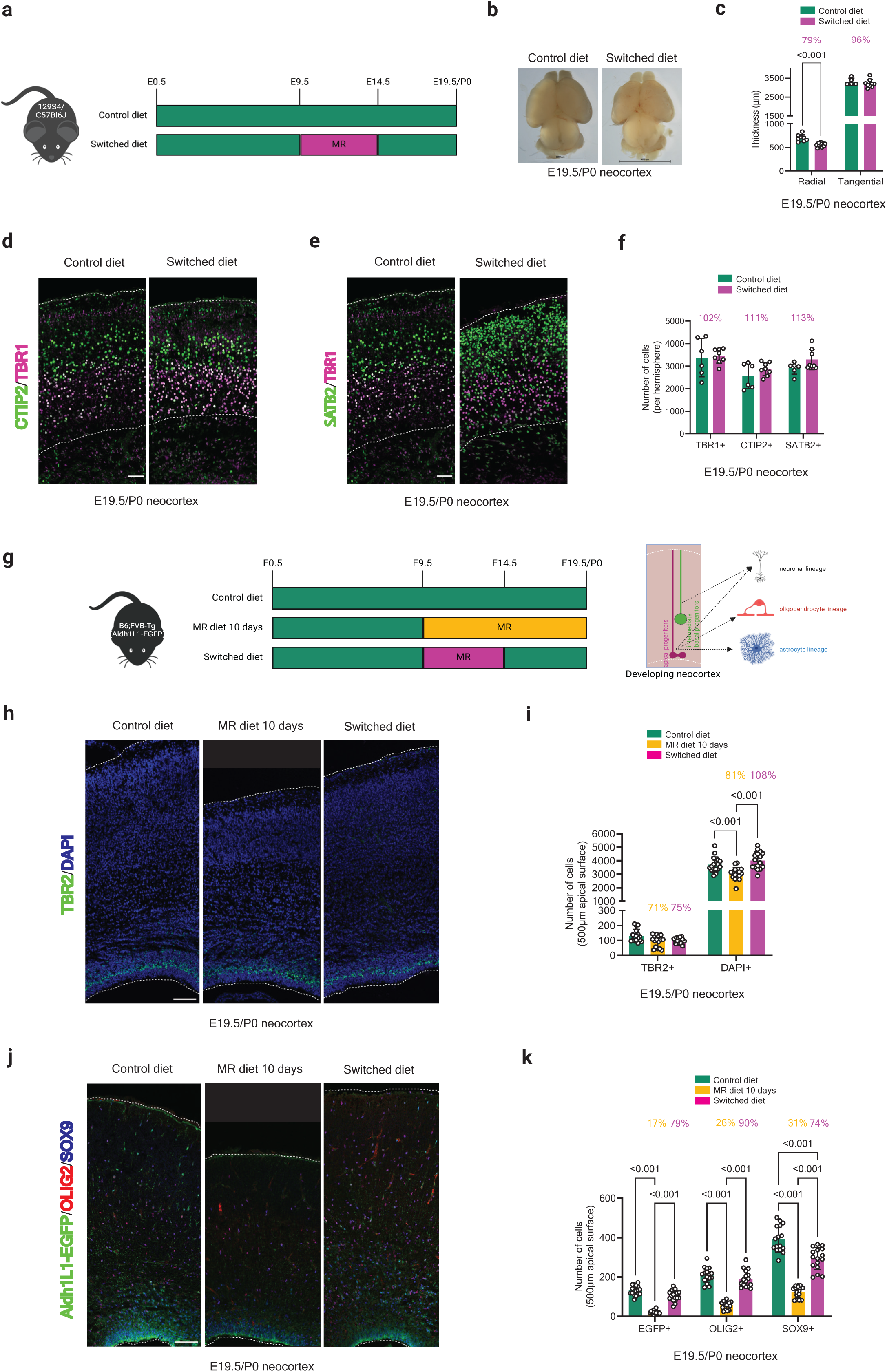
Growth defect, including neuronal production, is reversible in favorable nutrition conditions. **a**, Schematic of study design representing dietary switch regimen. **b,** Dorsal view of fetal brain at E19.5/P0 from pups exposed to control diet or switched diet. Scale bars: 5000 μm. **c,** Quantification of radial length and tangential perimeter of E19.5/P0 neocortex from pups exposed to control and switched diet (n= 6-7 embryos). **d-f,** Representative images and quantification of E19.5/P0 neocortex coronal sections immunostained for TBR1, CTIP2, and SATB2, in condition of control diet or switched diet (n= 6-7 embryos). Data are mean±s.d. Statistical analysis was carried out using multiple unpaired t test with Holm-Šídák correction. **g,** Schematic of study design representing dietary MR and switch regimen as well as illustrating the production of glial cells. Created with BioRender.com. **h-i,** Representative images and quantification of E19.5/P0 neocortex coronal sections immunostained for TBR2 in condition of control diet, 10 days of MR or switched diet (n= 4-5 embryos). Nuclei were stained with DAPI. **j-k,** Representative images and quantification of E19.5/P0 neocortex coronal sections from Aldh1L1-EGFP embryos immunostained for OLIG2 and SOX9, in condition of control diet, 10 days of MR or switched diet (n= 4-5 embryos). Scale bars: 100 μm. Data are mean±s.d. Statistical analysis was carried out using multiple unpaired t test with Holm-Šídák correction or two-way ANOVA with Tukey’s multiple comparision. p-values are indicated when differences are statistically significant. The percentage values at the top of the graph indicate the magnitude of the mean difference between conditions.

**Figure 4.**
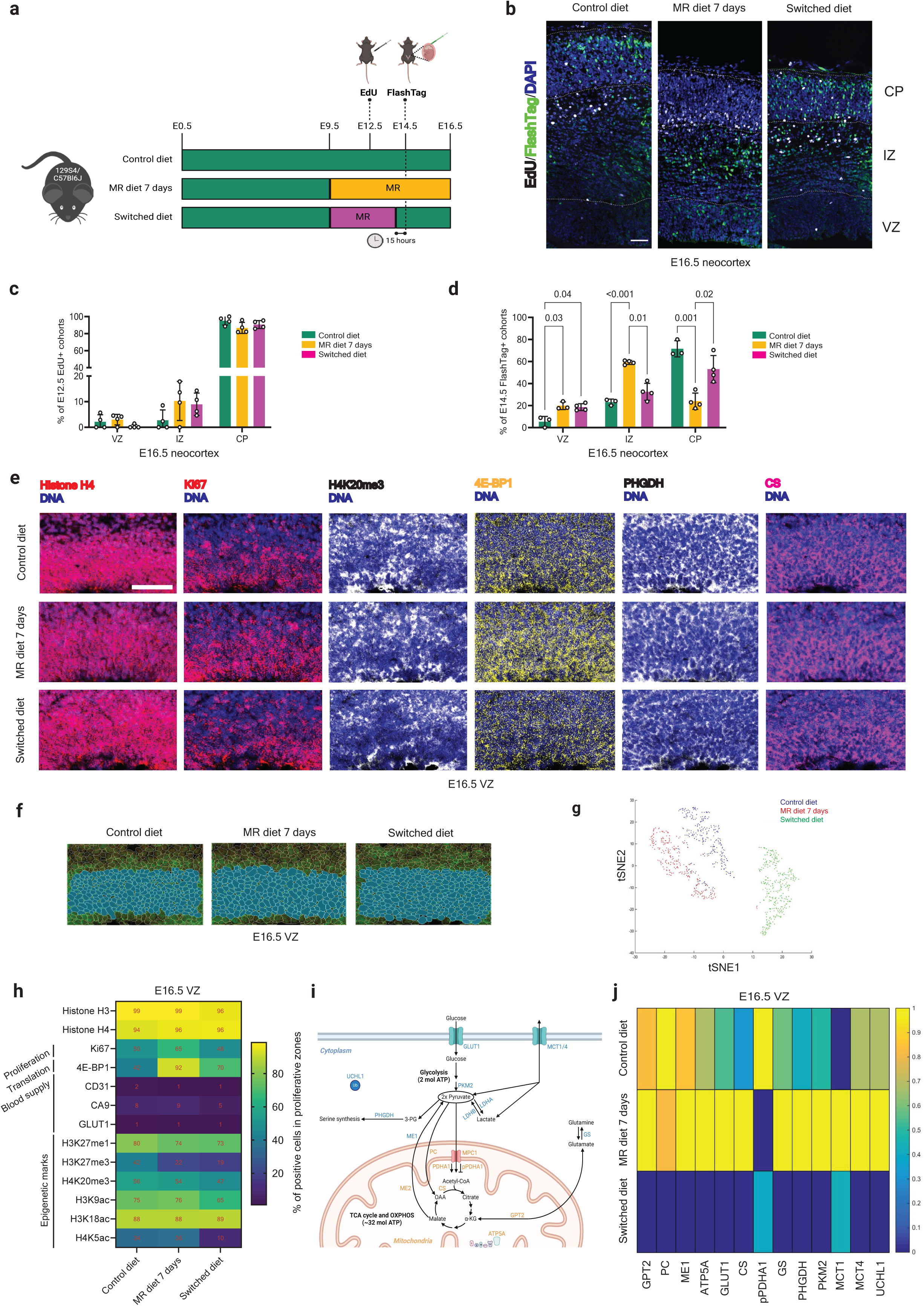
Progenitor behavior and molecular profiles in MR and catch-up growth conditions. **a**, Schematic of study design coupling birthdating strategies with dietary conditions. Created with BioRender.com. **b,** Representative images of coronal sections of E16.5 embryos exposed to the different dietary regimen, showing EdU and FlashTag-labelled cohort of cells born respectively at E12.5 and E14.5. Nuclei were stained with DAPI. **c,** Quantification of EdU+ cells in the ventricular zone (VZ), intermediate zone (IZ), and cortical plate (CP) in the different dietary regimens (n= 4 embryos). **d,** Quantification of FlashTag+ cells in the ventricular zone (VZ), intermediate zone (IZ), and cortical plate (CP) in the different dietary regimens (n= 4 embryos). Scale bars: 50 μm. Data are mean±s.d. Statistical analysis was carried out using two-way ANOVA with Tukey’s multiple comparision. p-values are indicated when differences are statistically significant. The percentage values at the top of the graph indicate the magnitude of the mean difference between conditions. **e,** Reconstituted images of the VZ from spatial IMC data for selected markers, in the three dietary conditions. Scale bars: 50 μm. **f,** Segmentation of IMC images in the three diet conditions (top). **g,** t-SNE unsupervised clustering of IMC data. **h,** Heatmaps representation of marker-specific cell frequencies in the three dietary conditions. The numbers represent the fraction of nuclei positive for each marker in each dietary condition. **i,** Simplified schematic representation of energy metabolism highlighting tested cytoplasmic metabolic markers in blue and tested mitochondrial metabolic markers in orange. **j,** Heatmaps representation of mean intensity signal for selected metabolic markers in the three dietary conditions.

## Supporting information

Materials and reagents

Diet composition

List of IMC antibodies

Quantification

## ACKNOWLEDGEMENTS

We acknowledge the help and contribution of the ANEXPLO mouse facility and the TRI imaging platforms at the CBI. Brice Ronsin and Christian Rouvière provided custom scripts for semi-automated counting. We acknowledge the SIMCaT (Spatial Imaging Mass Cytometry and Transcriptomics) facility of Montpellier Cancer Research Institute for technical and scientific expertise. This work was funded by Agence Française pour la Recherche (ANR-21-CE16-0024-01). Both CNRS and Université de Toulouse provided core funding. Sulov SAHA was the recipient of a 3-year PhD scholarship from the French Ministry of Research.

## AUTHOR INFORMATION

### Authors and Affiliations

Molecular, Cellular and Developmental Biology Unit, UMR 5077, Center for Integrative Biology, CNRS, Université Paul Sabatier, Toulouse, France

Sulov Saha, Clémence Debacq, Christophe Audouard, Thomas Jungas, Mohamad Ali Fawal, David Ohayon, Alice Davy

Institut de Recherche en Cancérologie de Montpellier (IRCM), INSERM U1194; Université de Montpellier, Institut régional du Cancer de Montpellier (ICM), Montpellier, France

Pierrick Dupré, Henri-Alexandre Michaud, Matthieu Lacroix, Laurent Le Cam

### Contributions

SS performed experiments, analyzed and interpreted data, wrote a draft and edited the manuscript; CA, TJ, CD and CC performed experiments, analyzed and interpreted data; MAF participated in the conceptualization of the study and edited the manuscript; HAM and PD performed the IMC experiments and processed data; ML and LL participated in the conception of IMC experiments, analyzed and interpreted IMC experiments and edited the manuscript; DO performed experiments, performed bioinformatics analyses, interpreted data and edited the manuscript; AD conceptualized and managed the study, analyzed and interpreted data, wrote the manuscript.

### Corresponding author

Correspondence to Alice Davy

## REFERENCES

1. Ozawa, H. et al. Transient Methionine Deprivation Triggers Histone Modification and Potentiates Differentiation of Induced Pluripotent Stem Cells. Stem Cells 41, 271–286 (2023).

2. Shiraki, N. et al. Methionine metabolism regulates maintenance and differentiation of human pluripotent stem cells. Cell Metab 19, 780–94 (2014).

3. Tang, S. et al. Methionine metabolism is essential for SIRT1-regulated mouse embryonic stem cell maintenance and embryonic development. EMBO J 36, 3175–3193 (2017).

4. Zhu, Q. et al. O-GlcNAcylation regulates the methionine cycle to promote pluripotency of stem cells. Proceedings of the National Academy of Sciences 117, 7755–7763 (2020).

5. Sanderson, S. M., Gao, X., Dai, Z. & Locasale, J. W. Methionine metabolism in health and cancer: a nexus of diet and precision medicine. Nat Rev Cancer 19, 625–637 (2019).

6. Korsmo, H. W. & Jiang, X. One Carbon Metabolism and Early Development: A Diet-Dependent Destiny. Trends Endocrinol Metab 32, 579–593 (2021).

7. Konycheva, G. et al. Dietary methyl donor deficiency during pregnancy in rats shapes learning and anxiety in offspring. Nutrition Research 31, 790–804 (2011).

8. Saha, S. et al. Dihydrofolate reductase activity controls neurogenic transitions in the developing neocortex. Development 150, dev201696 (2023).

9. Lauinger, L. & Kaiser, P. Sensing and Signaling of Methionine Metabolism. Metabolites 11, 83 (2021).

10. Mentch, S. J. et al. Histone Methylation Dynamics and Gene Regulation Occur through the Sensing of One-Carbon Metabolism. Cell Metab 22, 861–73 (2015).

11. Lin, D.-W., Chung, B. P. & Kaiser, P. S-adenosylmethionine limitation induces p38 mitogen-activated protein kinase and triggers cell cycle arrest in G1. J Cell Sci 127, 50–59 (2014).

12. Rakic, P. Mode of cell migration to the superficial layers of fetal monkey neocortex. Journal of Comparative Neurology 145, 61–83 (1972).

13. Gotz, M. & Huttner, W. B. The cell biology of neurogenesis. Nat. Rev. Mol. Cell Biol. 6, 777– 788 (2005).

14. Sessa, A., Mao, C. A., Hadjantonakis, A. K., Klein, W. H. & Broccoli, V. Tbr2 directs conversion of radial glia into basal precursors and guides neuronal amplification by indirect neurogenesis in the developing neocortex. Neuron 60, 56–69 (2008).

15. Bayraktar, O. A., Fuentealba, L. C., Alvarez-Buylla, A. & Rowitch, D. H. Astrocyte Development and Heterogeneity. Cold Spring Harb Perspect Biol 7, a020362 (2015).

16. Hippenmeyer, S. Principles of neural stem cell lineage progression: Insights from developing cerebral cortex. Current Opinion in Neurobiology 79, 102695 (2023).

17. Freret-Hodara, B. et al. Enhanced Abventricular Proliferation Compensates Cell Death in the Embryonic Cerebral Cortex. Cereb Cortex 27, 4701–4718 (2017).

18. Kischel, A., Audouard, C., Fawal, M.-A. & Davy, A. Ephrin-B2 paces neuronal production in the developing neocortex. BMC Dev Biol 20, 12 (2020).

19. Toma, K., Kumamoto, T. & Hanashima, C. The timing of upper-layer neurogenesis is conferred by sequential derepression and negative feedback from deep-layer neurons. J Neurosci. 34, 13259– 13276 (2014).

20. Govindan, S., Oberst, P. & Jabaudon, D. In vivo pulse labeling of isochronic cohorts of cells in the central nervous system using FlashTag. Nat Protoc 13, 2297–2311 (2018).

21. Salic, A. & Mitchison, T. J. A chemical method for fast and sensitive detection of DNA synthesis in vivo. Proceedings of the National Academy of Sciences 105, 2415–2420 (2008).

22. Du, Y., Gupta, P., Qin, S. & Sieber, M. The role of metabolism in cellular quiescence. Journal of Cell Science 136, jcs260787 (2023).

23. Scandella, V., Petrelli, F., Moore, D. L., Braun, S. M. G. & Knobloch, M. Neural stem cell metabolism revisited: a critical role for mitochondria. Trends in Endocrinology & Metabolism 34, 446– 461 (2023).

24. Jackson, B. T. & Finley, L. W. S. Metabolic regulation of the hallmarks of stem cell biology. Cell Stem Cell 31, 161–180 (2024).

25. Chang, Q. et al. Imaging Mass Cytometry. Cytometry Part A 91, 160–169 (2017).

26. Baharlou, H., Canete, N. P., Cunningham, A. L., Harman, A. N. & Patrick, E. Mass Cytometry Imaging for the Study of Human Diseases—Applications and Data Analysis Strategies. Front. Immunol. 10, (2019).

27. Dong, X. et al. Metabolic lactate production coordinates vasculature development and progenitor behavior in the developing mouse neocortex. Nat Neurosci 25, 865–875 (2022).

28. Schapiro, D. et al. histoCAT: analysis of cell phenotypes and interactions in multiplex image cytometry data. Nat Methods 14, 873–876 (2017).

29. Bankhead, P. et al. QuPath: Open source software for digital pathology image analysis. Sci Rep 7, 16878 (2017).

30. Alexander, M. P. et al. Exploring the single-cell immune landscape of kidney allograft inflammation using imaging mass cytometry. American Journal of Transplantation 24, 549–563 (2024).

31. Lu, S. & Epner, D. E. Molecular mechanisms of cell cycle block by methionine restriction in human prostate cancer cells. Nutr Cancer 38, 123–130 (2000).

32. Kaiser, P. Methionine Dependence of Cancer. Biomolecules 10, 568 (2020).

33. Otsuki, L. & Brand, A. H. Cell cycle heterogeneity directs the timing of neural stem cell activation from quiescence. Science 360, 99–102 (2018).

34. Nguyen, P. D. et al. Muscle Stem Cells Undergo Extensive Clonal Drift during Tissue Growth via Meox1-Mediated Induction of G2 Cell-Cycle Arrest. Cell Stem Cell 21, 107–119.e6 (2017).

35. Buzgariu, W., Crescenzi, M. & Galliot, B. Robust G2 pausing of adult stem cells in *Hydra*. Differentiation 87, 83–99 (2014).

36. Petrelli, F. et al. Mitochondrial pyruvate metabolism regulates the activation of quiescent adult neural stem cells. Sci Adv 9, eadd5220 (2023).

37. Kang, S., Antoniewicz, M. R. & Hay, N. Metabolic and transcriptomic reprogramming during contact inhibition-induced quiescence is mediated by YAP-dependent and YAP-independent mechanisms. Nat Commun 15, 6777 (2024).

38. Kagan, B. J. & Rosello-Diez, A. Integrating levels of bone growth control: From stem cells to body proportions. WIREs Developmental Biology 10, e384 (2021).

39. Tanner, J. M. Regulation of Growth in Size in Mammals. Nature 199, 845–850 (1963).

40. Gafni, R. I. & Baron, J. Catch-up growth: possible mechanisms. Pediatr Nephrol 14, 616–619 (2000).

41. Heintz, N. Gene Expression Nervous System Atlas (GENSAT). Nat Neurosci 7, 483–483 (2004).

42. Ohayon, D. et al. Sulfatase 2 promotes generation of a spinal cord astrocyte subtype that stands out through the expression of Olig2. Glia 67, 1478–1495 (2019).

43. Fawal, M. A. et al. Cross Talk between One-Carbon Metabolism, Eph Signaling, and Histone Methylation Promotes Neural Stem Cell Differentiation. Cell Rep 23, 2864–2873 e7 (2018).

44. Windhager, J. et al. An end-to-end workflow for multiplexed image processing and analysis. Nat Protoc 18, 3565–3613 (2023).

45. Jungas, T., Joseph, M., Fawal, M.-A. & Davy, A. Population Dynamics and Neuronal Polyploidy in the Developing Neocortex. Cereb Cortex Commun 1, tgaa063 (2020).

46. Darzynkiewicz, Z., Halicka, H. D. & Zhao, H. Analysis of Cellular DNA Content by Flow and Laser Scanning Cytometry. Adv Exp Med Biol 676, 137–147 (2010).

