## Supplementary figures for "Acute dietary methionine restriction highlights sensitivity of neocortex development to metabolic variations"

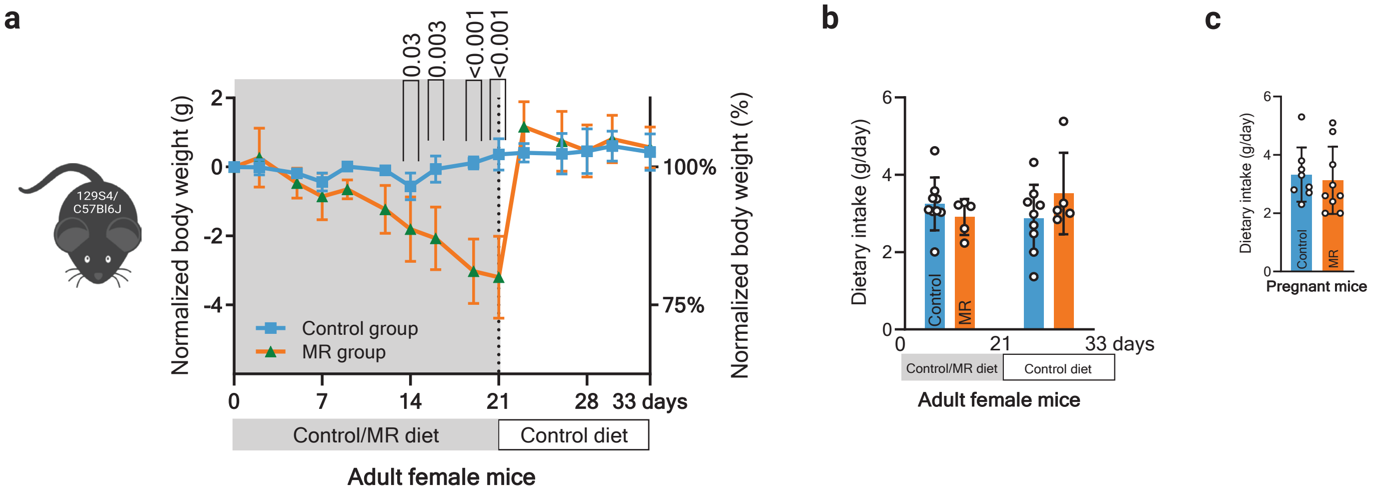


Figure S1. Food intake and body weight under MR diet. **a,** Trajectories of normalized body weight of adult female mice on control (SAFE #A04) and MR (SAFE AIN 2g Choline #U8958 Version 330) diet (n= 3 mice per group). **b,** Measurement of daily dietary intake of adult female mice over a 5-week period on control or MR diet (n= 3 mice per group). **c,** Measurement of daily dietary intake of adult females and pregnant mice over pregnancy on control or MR diet (n= 3 mice per group). Data are mean±s.d. Statistical analysis was carried out using mixed-effect analysis with Šídák multiple comparision or multiple Mann-Whitney test with Holm-Šídák correction. p-values are indicated when differences are statistically significant.

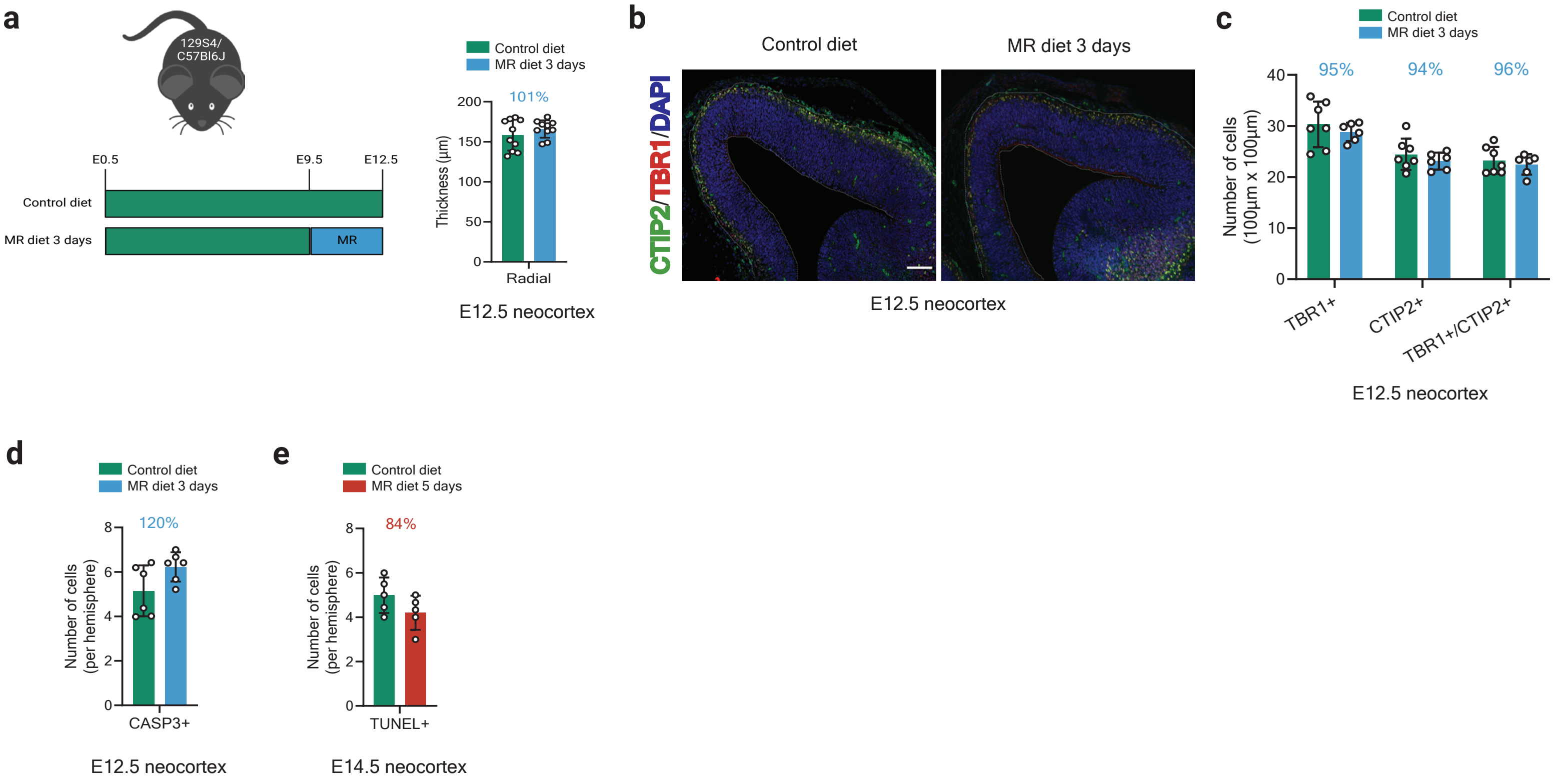


Figure S2. Onset of neurogenesis is maintained with no changes in cell death under methionine restriction. **a,** Schematic representation of the study design and quantification of radial thickness of neocortex of E12.5 embryo subjected to control or MR diet for 3 days. **b-c,** Representative images and quantification of E12.5 neocortex coronal sections immunostained for TBR1, CTIP2, and SATB2 in condition of control diet or 3-days MR (n= 4-5 embryos). Scale bars: 50 µm. **d-e,** Quantification of cell death by cleaved caspase-3 (CASP3) immunostaining in E12.5 and TUNEL assay in E14.5 neocortex coronal sections of embryos exposed to control diet or 3-days MR (n=3-4 embryos). Data are mean±s.d. Statistical analysis was carried out using multiple unpaired t-test with Holm-Šídák correction. p-values are indicated when differences are statistically significant.

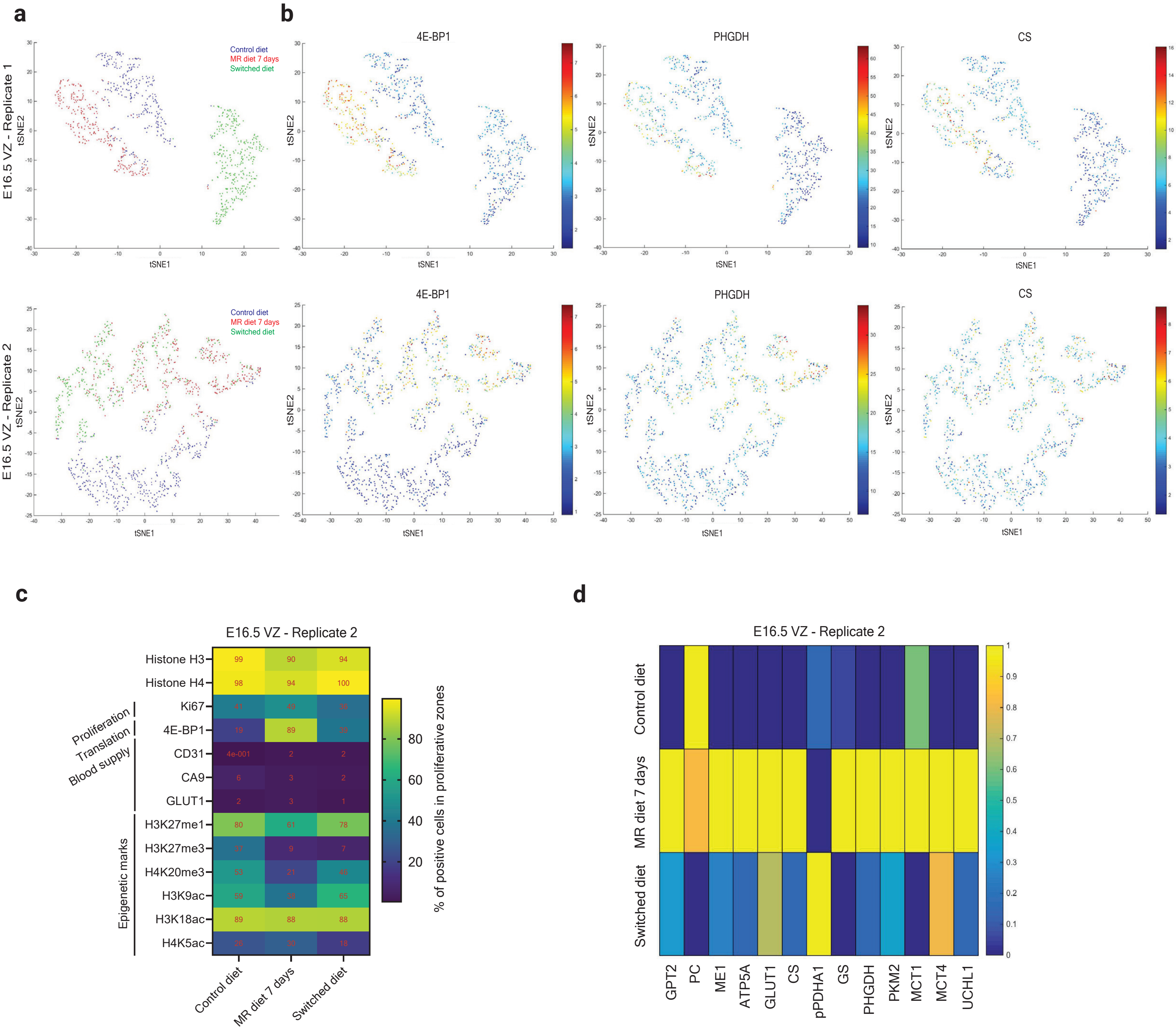


Figure S3. Bioinformatic analysis of IMC data. **a,** t-SNE unsupervised clustering of IMC data from two sets of samples (independent biological replicates). **b,** Clustered signal intensity for selected markers in the two replicates. **c,** Histogram representation of quantified IMC data for some panel markers in the three dietary conditions. The numbers represent the fraction of cells positive for each marker in each dietary condition, for the second replicate. **d,** Histogram representation of signal intensity for selected metabolic markers in the three diet conditions, for the second replicate.
