## Supplementary material for "Acute dietary methionine restriction highlights sensitivity of neocortex development to metabolic variations": Materials and reagents

**EXTENDED DATA- MATERIALS**

| REAGENT or RESOURCE | SOURCE | IDENTIFIER |
| --- | --- | --- |
| ***Antibodies*** | | |
| Mouse anti-PAX6 [1:100] | GeneTex | Cat#GTX34955<br>RRID:AB_2916297 |
| Rat anti-TBR2 (Eomes) [1:100] | Invitrogen | Cat#14-4875-82<br>RRID:AB_11042577 |
| Rabbit anti-TBR1 [1:100] | Abcam | Cat#ab31940<br>RRID:AB_2200219 |
| Rat anti-CTIP2 (Bcl11b) [1:300] | Abcam | Cat#ab18465<br>RRID:AB_2064130 |
| Mouse anti-SATB2 [1:50] | Abcam | Cat#ab51502<br>RRID:AB_882455 |
| Rabbit anti-pH3<br>(phosphorylated at ser10) [1:400] | Merck-Millipore | Cat#06-570<br>RRID:AB_310177 |
| Rabbit anti-CASP3<br>(cleaved at asp175) [1:400] | Cell Signaling Technology | Cat#9661<br>RRID:AB_2341188 |
| Chicken anti-GFP [1:500] | Abcam | #CatAB13970<br>RRID:AB_300798 |
| Rabbit anti-SOX9 [1:750] | Merck-Millipore | Cat#AB5535<br>RRID:AB_2239761 |
| Rabbit anti-OLIG2 [1:500] | Merck-Millipore | Cat#AB9610<br>RRID:AB_570666 |
| Mouse anti-OLIG2 [1:500] | Sigma-Aldrich | Cat#MABN50<br>RRID:AB_10807410 |
| Donkey anti-rabbit IgG AlexaFluor-488 conjugated [1:500] | ThermoFisher | Cat#A-21206<br>RRID:AB_2535792 |
| Donkey anti-rabbit IgG AlexaFluor-555 conjugated [1:500] | ThermoFisher | Cat#A-31572<br>RRID:AB_162543 |
| Donkey anti-rabbit IgG AlexaFluor-647 conjugated [1:500] | ThermoFisher | Cat#A-31573<br>RRID:AB_2536183 |
| Donkey anti-mouse IgG AlexaFluor-488 conjugated [1:500] | ThermoFisher | Cat#R37114<br>RRID:AB_2556542 |
| Donkey anti-mouse IgG AlexaFluor-555 conjugated [1:500] | ThermoFisher | Cat#A-31570<br>RRID:AB_2536180 |
| Donkey anti-mouse IgG AlexaFluor-647 conjugated [1:500] | ThermoFisher | Cat#A-31571<br>RRID:AB_162542 |
| Donkey anti-rat IgG AlexaFluor-488 conjugated [1:500] | ThermoFisher | Cat#A-21208<br>RRID:AB_141709 |
| Donkey Anti-Chicken IgY H&L (FITC) | Abcam | #CatAB63507<br>RRID: AB_1139472 |
| DAPI [1:1000] | ThermoFisher | Cat#62248 |
| ***Oligonucleotides*** | | |
| Genotyping primer for GFP-Fw<br>5'- CGCACCATCTTCTTCAAGGACGAC-3′ | Ohayon *et al.*, 2019 Glia | N/A |
| Genotyping primer for GFP-Rev 5′ -<br>AACTCCAGCAGGACCATGTGATCG-3′ | Ohayon *et al.*, 2019 Glia | N/A |
| ***Chemicals, Peptides, and Recombinant Proteins*** | | |
| BSA | Euromedex | Cat#04100811C |
| BSA Cohn fraction 30% | Sigma-Aldrich | Cat#126621 |
| Dako target retrieval solution pH9 | Agilent | Cat#S2367 |
| DMSO | Euromedex | Cat#UD8050-05 |

| | | |
|---|---|---|
| dPBS | ThermoFisher | Cat#14190094 |
| Ethanol | VWR chemicals | Cat#20821.330 |
| FBS (PBTA solution) | ThermoFisher | Cat#10500064 |
| Flashtag/CFSE | STEMCELL Technologies | Cat#75003 |
| HBSS | ThermoFisher | Cat#14025092 |
| Intercalator-Iridium | Standard Biotools | Cat#201192A |
| Mouse FcR blocking reagent | Miltenyi | Cat#130-092-575 |
| Mowiol | Sigma-Aldrich | Cat#81381 |
| Nuclease-free water | Sigma-Aldrich | Cat#W4502 |
| OCT media | Sakura | Cat#4583 |
| Paraformaldehyde | Fisher Scientific | Cat#15710 |
| PBS | Sigma-Aldrich | Cat#D1408 |
| Sucrose | Sigma-Aldrich | Cat#S0389 |
| SuperBlock blocking buffer | ThermoFisher | Cat#37536 |
| Taq 2x PCR MasterMix | abm | Cat#G888 |
| Triton X-100 | Sigma-Aldrich | Cat#T9284 |
| Triton X-100 | VWR | Cat#28817.295 |
| Tween 20 | Euromedex | Cat#2001-A |
| ***Critical Commercial Assays*** | | |
| Click-iT EdU cell proliferation kit | ThermoFisher | Cat#C10340 |
| DNeasy blood & tissue kit | Qiagen | Cat#69504 |
| MaxPar X8 multimetal labeling kit | Standard BioTools | Cat#201149A |
| Methionine assay kit (fluorometric) | Abcam | Cat#ab234041 |
| MethylFlash methylated DNA 5-mC quantification kit (colorimetric) | Epigentek | Cat#P-1034-96 |
| ***Experimental Models : Organisms/Strains*** | | |
| Wild type mice (129S4/C57Bl6J) | In-house breeding | N/A |
| Transgenic mice (Aldh1L1-EGFP/Rpl10a) JD130Htz | GENSAT, kindly provided by Dr. N. Rouach, Paris, FRANCE | RRID: IMSR_JAX:030247 |
| ***Experimental Diets*** | | |
| Control diet | SAFE | Cat#AO4 |
| Methionine-restricted (MR) diet | SAFE | Cat#U8958 Version 330 |
| ***Software and Algorithms*** | | |
| Adobe Illustrator | Adobe | RRID:SCR_010279 |
| Affinity Designer | Serif | RRID:SCR_016952 |
| BioRender | BioRender | RRID:SCR_018361 |
| CytExpert | Beckman Coulter | RRID:SCR_017217 |
| HistoCAT | Schapiro et al., 2017 Nat Methods | N/A |
| ImageJ | NIH | RRID:SCR_003070 |
| MCD Viewer | Standard BioTools | RRID:SCR_023007 |
| Prism10 | Graphpad | RRID:SCR_002798 |
| QuPath 0.5.1 | QuPath | RRID:SCR_018257 |
| Steinbock | Windhager et al., 2023 Nat Protoc | N/A |
| ***Other*** | | |
| 10kD spin column | Abcam | Cat#ab93349 |
| Analytical balance | Sartorius | Cat#A-120S |

| | | |
|---|---|---|
| Bench top centrifuge | Eppendorf | Cat#5415R |
| Black 96-well microplate | ThermoFisher | Cat#237105 |
| Centrifuge | Eppendorf | Cat#ST40R |
| Coverslips | Menzel-Gläser | Cat#17234914 |
| Cryostat CM1950 | Leica Biosystems | RRID:SCR_018061 |
| Cytoflex-S flow cytometer | Beckman Coulter | N/A |
| Eclipse 80i fluorescence microscope | Nikon | RRID:SCR_015572 |
| Eclipse TS100 inverted microscope | Nikon | RRID:SCR_020324 |
| Hyperion imaging system | Standard BioTools | Cat#400323 |
| Microtome | Microm | Cat#HM355 |
| Multiskan GO | ThermoFisher | Cat#N10588 |
| Nanodrop | ThermoFisher | Cat#ND-2000 |
| SMZ18 stereomicroscope | Nikon | N/A |
| SP8 inverted scanning confocal microscope | Leica Biosystems | N/A |
| SP8 upright scanning confocal microscope | Leica Biosystems | N/A |
| Spin tissue processor | Myr | Cat#STP120 |
| Steamer | Phillips | Cat#HD9104/01 |
| Superfrost plus slides | Thermoscientific | Cat#J1800AMNT |
| Varioskan flash | ThermoFisher | Cat#5250030 |
| Veriti thermal cycler | ThermoFisher | Cat#4375305 |
| Vetflurane | Vibrac | Cat#Vnr137317 |
| Vibratome | Leica Biosystems | Cat#VT1000S |
