## Supplementary material for "Acute dietary methionine restriction highlights sensitivity of neocortex development to metabolic variations": Diet composition

**S1 – Diet composition**

| Control diet (SAFE #AO4) | Amount (mg per kg) |
|---|---|
| **Amino acids** | |
| Arginine | 9000 |
| Cystine | 2500 |
| Lysine | 7200 |
| Methionine | 2800 |
| Tryptophan | 1900 |
| Glycine | 8100 |
| **Fatty acids** | |
| Palmitic acid | 5900 |
| Stearic acid | 600 |
| Palmitoleic acid | 150 |
| Oleic acid | 4800 |
| LA | 15000 |
| ALA | 1200 |
| **Minerals** | |
| Calcium | 7300 |
| Phosphorus | 5500 |
| Sodium | 2500 |
| Potassium | 6000 |
| Magnesium | 1600 |
| Manganese | 70 |
| Iron | 270 |
| Copper | 16 |
| Zinc | 55 |
| Chlorine | 4000 |
| **Vitamins** | |
| Vitamin A | 7500 IU |
| Vitamin D3 | 1000 IU |
| Vitamin E | 30 IU |
| Vitamin K3 | 2.5 |
| Vitamin B1 | 5 |
| Vitamin B2 | 6.5 |
| Vitamin B3 | 70 |
| Vitamin B5 | 10 |
| Vitamin B6 | 3 |
| Vitamin B9/Folate | 0.35 |
| Vitamin B12 | 0.01 |
| Biotin | 0.08 |
| Choline | 1600 |
| **Energy content** | **kcal per kg (%)** |
| Proteins | 644 (19.3) |
| Lipids | 279 (8.4) |
| Carbohydrates | 2416 (72.4) |

| MR diet (SAFE AIN 2g Choline #U8958 v330) | Amount (mg per kg) |
|---|---|
| **Amino acids** | |
| Arginine | 5880 |
| Cystine | 4116 |
| Lysine | 12696 |
| Methionine | 0 |
| Tryptophan | 2079 |
| Glycine | 2955 |
| **Fatty acids** | |
| Palmitic acid | 7420 |
| Stearic acid | 2555 |
| Palmitoleic acid | 350 |
| Oleic acid | 13020 |
| LA | 37030 |
| ALA | 5390 |
| Sum n-3 | 5390 |
| Sum n-6 | 37030 |
| Sum SFA | 10325 |
| Sum UFA | 55790 |
| Sum MUFA | 13370 |
| Sum PUFA | 42420 |
| Cholesterol | 1.1 |
| **Minerals** | |
| Calcium | 6333 |
| Phosphorus | 2678 |
| Sodium | 4574 |
| Potassium | 6323 |
| Magnesium | 823 |
| Manganese | 14 |
| Iron | 69 |
| Copper | 6.9 |
| Zinc | 45 |
| Chlorine | 4964 |
| **Vitamins** | |
| Vitamin A | 6437 IU |
| Vitamin D3 | 1625 IU |
| Vitamin E | 106 IU |
| Vitamin K3 | 8 |
| Vitamin B1 | 7.8 |
| Vitamin B2 | 7.5 |
| Vitamin B3 | 45 |
| Vitamin B5 | 20 |
| Vitamin B6 | 9.1 |
| Vitamin B9/Folate | 2.6 |
| Vitamin B12 | 0.033 |
| Biotin | 0.26 |
| Choline | 2062 |
| **Sugars** | |

| | |
|---|---|
| Glucose | <0.5 % |
| Sucrose | 13 % |
| **Energy content** | **kcal per kg (%)** |
| Proteins | 561.4 (14.6) |
| Lipids | 640.4 (16.6) |
| Carbohydrates | 2647.4 (68.8) |
