## Supplementary material for "Acute dietary methionine restriction highlights sensitivity of neocortex development to metabolic variations": List of IMC antibodies

**S2 – IMC antibodies**

| Antibody target | Metal isotopes | Species (type) | Source | Identifier | Discarded? |
|---|---|---|---|---|---|
| 4E-BP1 | 145Nd | Rabbit (mono) | Cell Signaling Technology | Cat#9644 | |
| ATP5A | 160Gd | Rabbit (mono) | Abcam | Cat#ab231692 | |
| CA9 | 147Sm | Goat (poly) | R&D System | Cat#AF2188 | |
| CASP3 | 172Yb | Rabbit (mono) | Standard Biotools | Cat#3172027D | |
| CD31 | 151Eu | Rabbit (mono) | Fluidigm | Cat#91H027151 | |
| CK8 | 113Cd | Mouse (mono) | BioLegend | Cat#904804 | |
| CS | 115In | Rabbit (mono) | Abcam | Cat#ab233838 | |
| ECADHERIN | 158Gd | Rabbit (mono) | Standard Biotools | Cat#3158029D | Yes |
| GLUT1 | 165Ho | Rabbit (mono) | Abcam | Cat#ab196357 | |
| GPT2 | 166Er | Rabbit (poly) | Proteintech | Cat#16757-I-AP | |
| GS | 154Sm | Rabbit (poly) | Proteintech | Cat#11037-2-AP | |
| H3K18ac | 170Er | Rabbit (mono) | Cell Signaling Technology | Cat#13998BF | |
| H3K27me1 | 161Dy | Mouse (mono) | Active Motif | Cat#61015 | |
| H3K27me3 | 155Gd | Mouse (mono) | Active Motif | Cat#61017 | |
| H3K4me3 | 141Pr | Rabbit (mono) | Cell Signaling Technology | Cat#9751 | Yes |
| H3K9ac | 148Nd | Rat (mono) | Active Motif | Cat#61663 | |
| H3K9me1 | 149Sm | Mouse (mono) | BioLegend | Cat#824201 | Yes |
| H3K9me2/3 | 142Nd | Mouse (mono) | Cell Signaling Technology | Cat#5327 | Yes |
| H4K20me1 | 144Nd | Mouse (mono) | BioLegend | Cat#828001 | |
| H4K20me3 | 150Nd | Mouse (mono) | BioLegend | Cat#827701 | |
| H4K5ac | 162Dy | Mouse (mono) | Active Motif | Cat#61523 | |
| Histone H3 | 111Cd | Rabbit (mono) | Cell Signaling Technology | Cat#4499S | |
| Histone H4 | 112Cd | Rabbit (mono) | Cell Signaling Technology | Cat#13919 | |
| Ki67 | 168Er | Mouse (mono) | Standard Biotools | Cat#3168022D | |
| LDHA | 156Gd | Rabbit (mono) | Abcam | Cat#ab219591 | |
| LDHB | 164Dy | Mouse (mono) | Ab frontier | Cat#YIF-LF-MA0436 | |
| MCT1 | 173Yb | Rabbit (poly) | Merck | Cat#AB3538P | |
| MCT4 | 174Yb | Rabbit (poly) | Proteintech | Cat#49309 | |
| ME1 | 153Eu | Mouse (mono) | GeneTex | Cat#GTX632190 | |
| ME2 | 171Yb | Rabbit (mono) | Abcam | Cat#ab240113 | |
| MPC1 | 146Nd | Rabbit (mono) | Cell Signaling Technology | Cat#14462BF | |
| pAKT | 152Sm | Rabbit (mono) | Cell Signaling Technology | Cat#4060BF | |
| PC | 167Er | Rabbit (poly) | Proteintech | Cat#16588-1-AP | |
| PDHA1 | 163Dy | Mouse (mono) | ThermoFisher | Cat#459400 | |
| PHGDH | 159Tb | Mouse (mono) | ThermoFisher | Cat#MA5-29479 | |
| PKM2 | 169Tm | Rabbit (mono) | Cell Signaling Technology | Cat#4053BF | |
| pPDHA1 | 175Lu | Rabbit (poly) | Abcam | Cat#ab92696 | |
| UCHL1 | 176Yb | Rabbit (mono) | Abcam | Cat#ab220823 | |
| VIMENTIN | 143Nd | Rabbit (mono) | Standard Biotools | Cat#3143027D | |
