## Supplementary material for "Acute dietary methionine restriction highlights sensitivity of neocortex development to metabolic variations": Quantification

**Body**

| Control diet | MR diet (10 days) |
|---|---|
| 1690 | 860 |
| 1520 | 1030 |
| 1440 | 790 |
| 1670 | 950 |
| | 760 |

**Figure 1C**

**measurement= wet mass (mg)**

**samples= E19.5/P0**

**number of samples= 4-5 animals per condition**

**different littermates used= 2**

**Brain**

| Control diet | MR diet (10 days) |
|---|---|
| 106 | 55 |
| 87 | 51 |
| 77 | 46 |
| 84 | 49 |
| | 45 |

**Liver**

| Control diet | MR diet (10 days) |
|---|---|
| 105 | 55 |
| 77 | 63 |
| 70 | 77 |
| 62 | 80 |
| | 59 |

**Heart**

| Control diet | MR diet (10 days) |
|---|---|
| 18 | 15 |
| 20 | 15 |
| 12 | 30 |
| 12 | 8 |
| | 18 |

**Figure 1D**

**measurement= wet mass (mg)**

**samples= E14.5**

**number of samples= 3-5 animals per condition**

**different littermates used= 2**

**Body**

| Control diet | MR diet (5 days) |
|---|---|
| 230 | 220 |
| 240 | 220 |
| 230 | 210 |
| 240 | 240 |
| 250 | |

**Brain**

| Control diet | MR diet (5 days) |
|---|---|
| 25 | 21 |
| 26 | 20 |
| 24 | 20 |
| 25 | 20 |
| 27 | |

**Liver**

| Control diet | MR diet (5 days) |
|---|---|
| 27.4 | 25.7 |
| 25.2 | 27.2 |
| 28.4 | 27.4 |
| 28.6 | |

**Heart**

| Control diet | MR diet (5 days) |
|---|---|
| 2.4 | 2.2 |
| 1.8 | 2 |
| 2.2 | 2.1 |
| 1.9 | |

**Neocortex**

| Control diet | MR diet |
|---|---|
| 96.3297605 | 87.90524515 |
| 101.910262 | 86.2932725 |
| 116.403421 | 84.8466363 |
| 99.8503991 | 61.154732 |
| 102.654732 | 85.6221209 |

**Liver**

| Control diet | MR diet |
|---|---|
| 113.469555 | 89.04435576 |
| 110.296237 | 92.249829 |
| 107.76374 | 97.4262258 |
| 105.303877 | 89.0443558 |
| 110.675257 | 92.249829 |

**Heart**

| Control diet | MR diet |
|---|---|
| 84.3425314 | 92.91470924 |
| 82.6605473 | 92.2620296 |
| 90.5558723 | 96.0265678 |
| 111.17081 | 83.7664766 |
| 79.2222349 | 87.9721779 |

**Figure 1E**

**measurement= methionine level (µM)**

**samples= E14.5**

**number of samples= 5 animals per condition**

**different littermates used= 2**

**experimental replicates= 2**

**Neocortex**

| Control diet | MR diet |
|---|---|
| 4.001370152 | 1.068718381 |
| 2.965851602 | 0.678225126 |
| 5.281408094 | 1.73693086 |
| 3.33315767 | 1.475021079 |

**Figure 1F**

**measurement= 5-mC level in total DNA (%)**

**samples= E14.5**

**number of samples= 4-5 animals per condition**

**different littermates used= 2**

**experimental replicates= 2**

**Liver**

| Control diet | MR diet |
|---|---|
| 1.807546374 | 1.365408938 |
| 1.262647555 | 2.244940978 |
| 2.654405565 | 2.750316189 |
| 1.47396712 | 2.532145868 |
| | 1.01918212 |

**Heart**

| Control diet | MR diet |
|---|---|
| 2.188026981 | 1.743254637 |
| 2.575885329 | 0.938026981 |
| 2.372470489 | 2.275505902 |
| 1.00916948 | 1.557230185 |

**Figure 1G**

**measurement= cells (%)**

**samples= E14.5**

**number of neocortex & liver samples= 5-7 animals per condition**

**number of heart samples= 3 (each pulldown of 4 animals) per condition**

**different litters used= 3**

**experimental replicates= 2**

**Neocortex**

| Cell cycle | Control diet | | | | | MR diet | | | | | |
|---|---|---|---|---|---|---|---|---|---|---|---|
| G0+G1 | 91.3 | 90.88 | 91.89 | 92.19 | 93.2 | 84.21 | 87.7 | 83.27 | 84.76 | 86.69 | 87.33 |
| S | 5.16 | 5.9 | 4.83 | 4.33 | 4.02 | 9.95 | 7.47 | 11.1 | 9.73 | 8.47 | 7.48 |
| G2+M | 3.48 | 2.99 | 3.11 | 3.37 | 2.6 | 5.62 | 4.27 | 5.42 | 5.31 | 4.52 | 5.28 |
| pH3 | 1.22 | 1.26 | 1.31 | 1.37 | 1.51 | 1.91 | 1.25 | 4.38 | 1.86 | 1.7 | 0.25 |

**Liver**

| Cell cycle | Control diet | | | | | MR diet | | | | | | |
|---|---|---|---|---|---|---|---|---|---|---|---|---|
| G0+G1 | 56.98 | 59.23 | 59.28 | 62.03 | 56.46 | 56.85 | 53.9 | 59.53 | 58.86 | 55.46 | 59.36 | 61.29 |
| S | 24.48 | 20.47 | 25.88 | 22.3 | 27.69 | 26.38 | 27.9 | 19.33 | 27.05 | 26.16 | 28.28 | 25.1 |
| G2+M | 18.64 | 19.03 | 14.56 | 14.27 | 15.51 | 16.07 | 17.9 | 18.95 | 14.86 | 17.11 | 11.37 | 14.11 |
| pH3 | 2.07 | 1.21 | 1.7 | 1.22 | 1.76 | 2.03 | 2 | 0.79 | 2.21 | 2.35 | 1.6 | 1.93 |

**Heart**

| Cell cycle | Control diet | | | MR diet | | |
|---|---|---|---|---|---|---|
| G0+G1 | 87.96 | 89.35 | 83.13 | 93.7 | 91.57 | 90.86 |
| S | 7.76 | 6.53 | 8.59 | 3.83 | 4.64 | 5.27 |
| G2+M | 3.87 | 4.06 | 8.12 | 1.83 | 3.47 | 3.88 |
| pH3 | 2.19 | 3.34 | 0.84 | 6.29 | 5.62 | 0.66 |

**Radial thickness**

| Control diet | MR diet (10 days) |
|---|---|
| 711 | 442 |
| 658 | 496 |
| 631 | 442 |
| 707 | 455 |
| 834 | 447 |
| 605 | 508 |
| 756 | 443 |
| 681 | 498 |
| | 488 |

**Figure 2C**

**measurement= thickness (µm)**

**samples= E19.5/P0 coronal sections**

**number of samples= 4-5 animals per condition**

**different litters used= 2**

**Tangential thickness**

| Control diet | MR diet (10 days) |
|---|---|
| 3603 | 2365 |
| 3189 | 1950 |
| 3201 | 2629 |
| 3194 | 2882 |
| 3407 | 2033 |
| | 2429 |
| | 1923 |
| | 2385 |
| | 2831 |

| Radial thickness | |
|---|---|
| **Control diet** | **MR diet (5 days)** |
| 334.70013 | 178.4543 |
| 385.595082 | 150.54 |
| 397.056036 | 173.5267 |
| 257.000522 | 169.0643 |
| 277.782808 | 205 |
| 365.324577 | 205 |
| 347.991315 | 223 |
| 405.913828 | 196 |
| 397.245671 | 245 |
| 317.16129 | |

**Figure 2D**

**measurement= thickness (µm)**

**samples= E14.5 head coronal sections**

**number of samples= 4-5 animals per condition**

**different litters used= 2**

**TBR1**

| Control diet | MR diet (10 days) |
|---|---|
| 2269 | 2215 |
| 2704 | 1615 |
| 4269 | 1343 |
| 3022 | 1922 |
| 4322 | 1723 |
| 3655 | 1300 |
| | 1459 |
| | 1629 |
| | 1366 |

**Figure 2H**

**measurement= IF counts (TBR1, CTIP2 & SATB2)**
**samples= E19.5/P0 coronal sections**
**number of samples= 3-5 animals per condition**
**total counted= 4 sections per hemisphere**
**different litters used= 2**
**datapoints= average per hemisphere**

**CTIP2**

| Control diet | MR diet (10 days) |
|---|---|
| 3005 | 1916 |
| 2064 | 1645 |
| 3084 | 1321 |
| 2164 | 1830 |
| 3154 | 1496 |
| 1934 | 1237 |
| | 1434 |
| | 1377 |
| | 1136 |

**SATB2**

| Control diet | MR diet (10 days) |
|---|---|
| 3192 | 2092 |
| 2421 | 1863 |
| 2983 | 1152 |
| 3031 | 1898 |
| 3011 | 1427 |
| 2813 | 1185 |
| | 1250 |
| | 1493 |
| | 1179 |

**TBR1**

| Control diet | MR diet (5 days) |
|---|---|
| 1310 | 568 |
| 1134 | 457 |
| 1229 | 460 |
| 1257 | 524 |
| 1197 | 859 |
| | 806 |
| | 888 |

**Figure 2L**

**measurement= IF counts (TBR1, CTIP2 & SATB2)**

**samples= E14.5 head coronal sections**

**number of samples= 3-5 animals per condition**

**total counted= 2 sections per hemisphere**

**different litters used= 2**

**datapoints= counts per hemisphere**

**CTIP2**

| Control diet | MR diet (5 days) |
|---|---|
| 1015 | 394 |
| 1010 | 431 |
| 1002 | 342 |
| 1101 | 363 |
| 1124 | 565 |
| | 707 |
| | 635 |

**SATB2**

| Control diet | MR diet (5 days) |
|---|---|
| 869 | 208 |
| 818 | 206 |
| 717 | 215 |
| 786 | 205 |
| 673 | 269 |
| | 326 |
| | 324 |

**PAX6**

| Control diet | MR diet (5 days) |
|---|---|
| 1965 | 1112 |
| 1996 | 1246 |
| 1939 | 1138 |
| 1837 | 1136 |
| 1953 | 1617 |
| | 1136 |
| | 1342 |

**Figure 2N**

**measurement= IF counts (PAX6 & TBR2)**
**samples= E14.5 head coronal sections**
**number of samples= 3-5 animals per condition**
**total counted= 2 sections per hemisphere**
**different litters used= 2**
**datapoints= counts per hemisphere**

**TBR2**

| Control diet | MR diet (5 days) |
|---|---|
| 1015 | 472 |
| 1048 | 593 |
| 1050 | 444 |
| 948 | 558 |
| 989 | 733 |
| | 597 |
| | 699 |

TBR2

| Control diet | MR diet (5 days) |
|---|---|
| 31.2987013 | 48.55875831 |
| 39.6258503 | 50.79726651 |
| 44.6071904 | 49.42528736 |
| 42.0413123 | 46.78362573 |
| 40.3162055 | 51.94508009 |
| 40.1818182 | 51.30434783 |
| | 44.58715596 |
| | 57.43944637 |

**Figure 2P**

**measurement= IF counts (EdU)**

**samples= E14.5 head coronal sections**

**number of samples= 3-4 animals per condition**

**total counted= 2 sections per hemisphere**

**datapoints= % of EdU+ cohorts per hemisphere**

pH3

| Control diet | MR diet (5 days) |
|---|---|
| 68.9189189 | 28.72340426 |
| 66.6666667 | 34.32835821 |
| 81.0126582 | 28.69565217 |
| 76.8518519 | 32.71028037 |
| 80.1980198 | 27.65957447 |
| 72.8571429 | 28.16901408 |
| | 33.33333333 |
| | 46.59090909 |

pH3

| Control diet | MR diet (5 days) |
|---|---|
| 21.8724559 | 32.45334547 |
| 23.8607822 | 34.71882641 |
| 28.5676463 | 30.44263776 |
| 29.0388029 | 30.97713098 |
| 29.2896175 | 33.18476499 |
| 28.9019964 | 33.20235756 |
| | 27.09677419 |
| | 33.55391203 |

**Radial thickness**

| Control diet | Switched diet |
| --- | --- |
| 711 | 479 |
| 658 | 590 |
| 631 | 496 |
| 707 | 551 |
| 834 | 542 |
| 605 | 615 |
| 756 | 580 |
| 681 | 596 |
| | 530 |
| | 578 |

**Figure 3C**

**measurement= thickness (µm)**

**samples= E19.5/P0 coronal sections**

**number of samples= 4-6 animals per condition**

**different litters used= 2**

**Tangential thickness**

| Control diet | Switched diet |
| --- | --- |
| 3603 | 2950 |
| 3189 | 3656 |
| 3201 | 3043 |
| 3194 | 3096 |
| 3407 | 3201 |
| | 3401 |
| | 3228 |
| | 3143 |
| | 3212 |

#### TBR1

| Control diet | Switched diet |
|---|---|
| 2269 | 3374 |
| 2704 | 3193 |
| 4269 | 2878 |
| 3022 | 3723 |
| 4322 | 3744 |
| 3655 | 3777 |
| | 3290 |
| | 3573 |

**Figure 3F**

**measurement= IF counts (TBR1, CTIP2 & SATB2)**
**samples= E19.5/P0 coronal sections**
**number of samples= 3-5 animals per condition**
**total counted= 4 sections per hemisphere**
**different litters used= 2**
**datapoints= average per hemisphere**

#### CTIP2

| Control diet | Switched diet |
|---|---|
| 3005 | 2877 |
| 2064 | 2565 |
| 3084 | 2415 |
| 2164 | 2746 |
| 3154 | 3066 |
| 1934 | 3184 |
| | 2823 |
| | 3230 |

#### SATB2

| Control diet | Switched diet |
|---|---|
| 3192 | 4116 |
| 2421 | 3749 |
| 2983 | 3042 |
| 3031 | 3538 |
| 3011 | 2941 |
| 2813 | 3001 |
| | 2929 |
| | 3015 |

**Figure 3I**

**measurement= IF counts (TBR2 & DAPI)**
**samples= E19.5/P0 coronal sections**
**number of samples= 4 animals per condition**
**total counted= 4 sections per animal**
**different litters used= 2**
**datapoints= raw counts per 500µm apical surface**

**TBR2**

| Control diet | MR diet (10 days) | Switched diet |
|---|---|---|
| 104 | 36 | 90 |
| 91 | 96 | 102 |
| 96 | 144 | 81 |
| 100 | 124 | 95 |
| 201 | 34 | 83 |
| 138 | 44 | 127 |
| 125 | 45 | 79 |
| 139 | 57 | 95 |
| 114 | 140 | 81 |
| 81 | 92 | 111 |
| 101 | 106 | 108 |
| 95 | 108 | 63 |
| 165 | 106 | 118 |
| 132 | 128 | 120 |
| 210 | 106 | 97 |
| 204 | 137 | 124 |

**DAPI**

| Control diet | MR diet (10 days) | Switched diet |
|---|---|---|
| 3380 | 2676 | 5129 |
| 3300 | 2998 | 4541 |
| 3044 | 2800 | 4696 |
| 2892 | 2965 | 4301 |
| 3731 | 3050 | 2877 |
| 3402 | 2619 | 3357 |
| 3377 | 2638 | 3557 |
| 3476 | 1928 | 4390 |
| 5113 | 3172 | 3339 |
| 4569 | 3136 | 3625 |
| 4183 | 2896 | 3555 |
| 3560 | 3374 | 3545 |
| 3591 | 3332 | 4859 |
| 3739 | 3832 | 4513 |
| 4061 | 3786 | 3855 |
| 3958 | 3174 | 4074 |

**Figure 3K**

**measurement= IF counts (Aldh1L1-EGFP, OLIG2 & SOX9)**
**samples= E19.5/P0 coronal sections**
**number of samples= 4 animals per condition**
**total counted= 4 sections per animal**
**different litters used= 2**
**datapoints= raw counts per 500µm apical surface**

**EGFP**

| Control diet | MR diet (10 days) | Switched diet |
|---|---|---|
| 113 | 8 | 131 |
| 172 | 19 | 114 |
| 118 | 14 | 95 |
| 141 | 21 | 121 |
| 130 | 33 | 81 |
| 110 | 22 | 106 |
| 85 | 37 | 111 |
| 134 | 24 | 94 |
| 102 | 20 | 94 |
| 114 | 25 | 91 |
| 161 | 33 | 50 |
| 136 | 44 | 64 |
| 153 | 16 | 117 |
| 157 | 16 | 155 |
| 162 | 22 | 122 |
| 118 | 19 | 139 |

**OLIG2**

| Control diet | MR diet (10 days) | Switched diet |
|---|---|---|
| 151 | 60 | 150 |
| 250 | 59 | 167 |
| 147 | 89 | 141 |
| 246 | 69 | 150 |
| 244 | 73 | 191 |
| 294 | 45 | 210 |
| 218 | 54 | 243 |
| 219 | 66 | 178 |
| 220 | 69 | 204 |
| 165 | 63 | 187 |
| 185 | 78 | 144 |
| 215 | 57 | 162 |
| 196 | 32 | 230 |
| 225 | 30 | 288 |
| 193 | 24 | 247 |
| 201 | 27 | 161 |

**SOX9**

| Control diet | MR diet (10 days) | Switched diet |
|---|---|---|
| 346 | 140 | 356 |

| | | |
|---|---|---|
| 461 | 108 | 301 |
| 284 | 130 | 250 |
| 486 | 122 | 198 |
| 474 | 157 | 342 |
| 394 | 144 | 326 |
| 349 | 144 | 273 |
| 355 | 156 | 210 |
| 458 | 83 | 290 |
| 358 | 80 | 366 |
| 404 | 113 | 202 |
| 324 | 157 | 310 |
| 502 | 82 | 293 |
| 409 | 82 | 360 |
| 336 | 152 | 337 |
| 338 | 144 | 259 |

### Figure 4C & 4D

**measurement= IF counts (EdU & FlashTag cohorts)**
**samples= E16.5 coronal sections**
**number of samples= 4-5 animals per condition**
**total counted= 2 sections per hemisphere**
**different litters used= 2**
**datapoints= fraction of cohorts (%)**

**EdU**

| Tissue regior | Control diet | MR diet (7 days) | MR diet (5 days) |
|---|---|---|---|
| VZ | 2.142857 | 2.984482 | 0.357143 |
| IZ | 2.738095 | 10.292731 | 8.966346 |
| CP | 95.119048 | 86.722787 | 90.676511 |

**FlashTag**

| | Control diet | MR diet (7 days) | MR diet (5 days) |
|---|---|---|---|
| VZ | 12.953543 | 22.039023 | 14.468697 |
| IZ | 23.488871 | 59.168384 | 34.969067 |
| CP | 63.557586 | 18.792593 | 50.562237 |

**<u>Supp Figure 1A</u>**

**measurement= normalized body weight (g)**

**samples= adult female mice**

**number of samples= 3-4 animals per condition**

| Day | Control diet | | | MR diet | | | |
|---|---|---|---|---|---|---|---|
| 0 | 20 | 20 | 20 | 20 | 20 | 20 | 20 |
| 2 | 19.79 | 19.96 | 20.22 | 19.81 | 19.77 | 19.959 | 21.55 |
| 5 | 19.92 | 19.64 | 19.9 | 19.19 | 19.2 | 19.629 | 20.1 |
| 7 | 19.43 | 19.4 | 19.87 | 18.97 | 18.91 | 18.575 | 20.13 |
| 9 | 20.16 | 19.89 | 19.97 | 19.34 | 19.48 | 18.97 | 19.609 |
| 12 | 20.02 | 19.73 | 19.96 | 18.41 | 19.59 | 18.007 | 19.066 |
| 14 | 19.3 | 19.13 | 19.88 | 18.39 | 18.85 | 16.829 | 18.723 |
| 16 | 20.05 | 19.51 | 20.26 | 18.15 | 18.32 | 16.598 | 18.637 |
| 19 | 20.3 | 19.96 | 20.09 | 17.17 | 17.33 | 15.626 | 17.77 |
| | | | | **Control diet** | | | |
| 21 | 20.87 | 20.21 | 20.01 | 17.38 | 17.08 | 15.066 | 17.685 |
| 23 | 20.67 | 20.43 | 20.13 | 21.71 | 20.48 | 20.593 | 21.868 |
| 26 | 20.75 | 20.69 | 19.71 | 21.38 | 20.21 | 19.802 | 21.582 |
| 28 | 21.02 | 20.6 | 19.75 | 21.11 | 19.98 | 19.654 | 21.105 |
| 30 | 20.51 | 21.07 | 20.21 | 20.86 | 20.44 | 20.189 | 21.763 |
| 33 | 20.57 | 20.87 | 19.85 | 20.64 | 20.37 | 19.909 | 21.334 |

**Supp Figure 1B**

**measurement= dietary intake (g/day)**

**samples= adult female mice**

**number of samples= 3 animals per condition**

| Day 0 | Control diet | MR diet |
|---|---|---|
| | 2.003333 | 3.29625 |
| | 3.586667 | 2.979167 |
| | 3.028333 | 3.50375 |
| | 3.375 | 4.31875 |
| | 3.388889 | 2.478333 |
| | 3.05 | 3.215 |
| | 3.068333 | 2 |
| | 4.618889 | 1.363333 |
| | 3.096667 | 2.6975 |
| **Day 21** | | **Control diet** |
| | 2.606667 | 5.38375 |
| | 2.232222 | 3.265833 |
| | 3.285 | 2.84125 |
| | 3.183333 | 3.005 |
| **Day 33** | 3.216667 | 3.0825 |

| Control diet | MR diet |
|---|---|
| 2.9 | 3.66666667 |
| 2.7 | 4.8 |
| 2.3 | 2.6 |
| 2.8 | 2.6 |
| 5.3 | 2.4 |
| 3.3 | 2 |
| 3.8 | 5.1 |
| 3.5 | 2 |
| | 2.96666667 |

**Supp Figure 1C**

**measurement= dietary intake (g/day)**

**samples= pregnant female mice (E9,5-16,5)**

**number of samples= 3 animals per condition**

**different litters used= 2**

**experimental replicates= 2**

| Radial thickness | |
|---|---|
| Control diet | MR diet (3 days) |
| 177.26999 | 175.3333333 |
| 180.39194 | 181 |
| 178.51877 | 173.3333333 |
| 165.576869 | 163.3333333 |
| 147.204667 | 164.3333333 |
| 151.972736 | 170.3333333 |
| 137.744213 | 171.6666667 |
| 175.661713 | 147 |
| 132.06794 | 164.6666667 |
| 136.079173 | 149 |

**Supp Figure 2A**

**measurement= thickness (µm)**

**samples= E12.5 head coronal sections**

**number of samples= 5 animals per condition**

**different litters used= 2**

**TBR1**

| Control diet | MR diet (3 days) |
|---|---|
| 35.77 | 26.1971219 |
| 29.49 | 27.2910083 |
| 34.64 | 30.4206684 |
| 25.08 | 28.6540101 |
| 24.48 | 29.8217743 |
| 29.56 | 30.6621509 |
| 33.19 | |

**Supp Figure 2B**

**measurement= IF counts (TBR1, CTIP2 & SATB2)**

**samples= E12.5 head coronal sections**

**number of samples= 4-5 animals per condition**

**total counted= 2 sections per hemisphere**

**different litters used= 2**

**datapoints= normalized per 100µm x 100µm area (lateral)**

**CTIP2**

| Control diet | MR diet (3 days) |
|---|---|
| 23.14 | 20.9831077 |
| 26.1 | 22.9819018 |
| 20.67 | 24.4070417 |
| 23.45 | 25.1506105 |
| 25.56 | 23.7650247 |
| 22.23 | 21.3932226 |
| 29.98 | |

**TBR1/CTIP2**

| Control diet | MR diet (3 days) |
|---|---|
| 21.04 | 19.1391099 |
| 25.8 | 24.4182706 |
| 24.68 | 23.2076525 |
| 20.82 | 23.6611259 |
| 22.23 | 23.3842899 |
| 27.26 | 20.9767642 |
| 20.9 | |

**<u>Supp Figure 2C</u>**

**measurement= IF counts (CASP3)**

**samples= E12.5 head coronal sections**

**number of samples= 4-5 animals**

**total counted= 4 sections per hemisphere**

**different litters used= 2**

**datapoints= average per hemisphere**

**CASP3**

| Control diet | MR diet |
|---|---|
| 6.42054575 | 5.66572238 |
| 4.01069519 | 6.39658849 |
| 5.91715976 | 6.41025641 |
| 4.3715847 | 6.66666667 |
| 6.19195046 | 7.00280112 |
| 3.98406375 | 5.21512386 |

**Supp Figure 2D**
**measurement= IF counts (TUNEL)**
**samples= E14.5 head coronal sections**
**number of samples= 5 animals**
**total counted= 4 sections per hemisphere**
**different litters used= 2**
**datapoints= average per animal**

**TUNEL**

| Control diet | MR diet |
|---|---|
| 5.5 | 5 |
| 5 | 4.666666667 |
| 6 | 3 |
| 4 | 4 |
| 4.446504878 | 4.333333333 |
